# Cholesterol Biosynthesis is a Targetable Vulnerability of *CEBPA*-mutant Acute Myeloid Leukemia

**DOI:** 10.64898/2026.02.20.706425

**Authors:** Sofie Ulfbeck Schovsbo, Yang Liu, Pedro Aragon-Fernandez, Sandra Gordon, Mikkel Bruhn Schuster, Jinyu Su, Sachin Pundhir, Nanna S Mikkelsen, Erwin M Schoof, Kim Theilgaard-Mönch, Kirsten Grønbæk, Rasmus O Bak, Bauke de Boer, Bo T Porse

## Abstract

Bi-allelic *CEBPA* mutations occur in 5-15% of acute myeloid leukemia (AML) patients. The precise molecular consequences of *CEBPA* mutations, especially in combination with frequently co-occurring mutations in *TET2*, *WT1*, and *GATA2,* remain incompletely understood. Here, we present a robust human model of *CEBPA*-mutant AML through gene editing of healthy bone marrow-derived hematopoietic stem cells. Loss of the CEBPA-p42 isoform expressed in healthy cells with concomitant upregulation of the leukemic CEBPA-p30 isoform resulted in a myeloproliferative phenotype. Concurrent loss-of-function mutations in *TET2* or *WT1* drove full leukemic transformation, while GATA2 haploinsufficiency promoted erythroid precursor accumulation without overt AML. Single-cell transcriptomics and low-input proteomics revealed enhanced myeloid output, increased interferon signaling and elevated cholesterol biosynthesis in leukemic cells. Targeting cholesterol synthesis enhanced chemosensitivity, highlighting a potential therapeutic vulnerability, particularly relevant for *CEBPA*-mutant patients harboring co-mutations in *TET2* or *WT1*, which have poor outcomes.

**Statement of significance:** Induction of CEBPA-p30 by CRISPR/Cas gene editing in healthy human BM HSCs drives overt AML *in vivo*. TET2 and WT1 loss accelerate leukemogenesis, while GATA2 haploinsufficiency redirects differentiation toward erythroid precursors potentially driving acute erythroid leukemia. CEBPA-p30 AML exhibits cholesterol biosynthesis dependency, revealing a therapeutic vulnerability to statins.

## Introduction

The transcription factor CCAAT enhancer binding protein alpha (CEBPA) is a master regulator of myeloid differentiation in healthy hematopoiesis and is mutated in approximately 5–15% of patients with acute myeloid leukemia (AML) (1,2). *CEBPA* is an intronless gene encoding several functional domains, including two N-terminal transactivation domains (TADs), required for transcriptional activation, and a C-terminal basic leucine zipper (bZIP) domain, essential for dimerization and DNA binding (3,4). Through alternative translation initiation, *CEBPA* gives rise to two protein isoforms: a full-length isoform (CEBPA-p42) and an N-terminally truncated isoform that lacks the first TAD (CEBPA-p30). Most *CEBPA*-mutant AML patients harbor biallelic mutations (*CEBPA^bi^*), typically consisting of an N-terminal frameshift or nonsense mutation (*CEBPA^N^*) on one allele and a C-terminal in-frame deletions/insertions (indels) or point mutations (*CEBPA^C^*) on the other allele (5). *CEBPA^N^* mutations abrogate expression of CEBPA-p42 and increase expression of CEBPA-p30, whereas *CEBPA^C^* mutations generate CEBPA-p42 variants that are unable to dimerize or bind DNA. Consequently, CEBPA-p30 homodimers constitute the only functional CEBPA entities in *CEBPA^bi^* AML, in contrast to normal hematopoietic cells where CEBPA-p42 predominates (3,4,6).

Insights into CEBPA-p30-driven AML have largely been derived from murine models. Whereas a full *Cebpa* knockout (KO) is incompatible with myeloid commitment and therefore does not induce leukemia, CEBPA-p30 alone is sufficient to drive myeloid commitment, and both *Cebpa^N/N^* and *Cebpa^N/C^* mice develop AML within one year, characterized by hyperproliferation of immature granulocytes (7–9). Mechanistically, AML arises from loss of CEBPA-p42 activity, which enhances hematopoietic stem cell (HSC) self-renewal and proliferation, combined with CEBPA-p30-specific transcriptional programs resulting in myeloid progenitor expansion. Disease maintenance relies on a subset of leukemic granulocyte-macrophage progenitors (8,9). Follow-up studies identified p30-specific targets including Musashi RNA-binding protein 2, which sustains leukemic cell proliferation (10), and CD73, which inhibits apoptosis and promotes cell proliferation (11). *CEBPA* mutations often co-occur with additional genetic lesions, including recurrent co-mutations in *GATA2*, *TET2*, *WT1, NRAS*, and *CSF3R* among others (12). Combinatorial mouse models have revealed co-operation between *Cebpa* mutations and mutations in *Tet2* (12), *Gata2* (13), *Csf3r* (14), and *Asxl1* (15) in regards to AML development.

In contrast, few studies in human cells have examined the molecular consequences of CEBPA-p30 induction and CEBPA-p42 loss, especially in the context of commonly co-occurring AML mutations. Transcriptional profiling of primary AML samples showed that *CEBPA^bi^* AML is characterized by a distinct gene regulatory network involving RUNX1 and AP-1 family transcription factors, which together execute the aberrant transcriptional programs orchestrated by CEBPA-p30 (16,17). Recently, an induced pluripotent stem cell model was developed combining *CEBPA^C^* mutations with *RUNX1/SRSF2,* but no human CEBPA-p30 model currently exists (18).

Studying the consequences of a single genetic mutation in primary AML patient material is challenged by both inter- and intra-patient heterogeneity, with multiple subclones often present within one patient (19). To overcome this challenge, we developed a robust human model of CEBPA-p30-driven leukemogenesis by CRISPR/Cas9-mediated gene editing of primary human bone marrow (BM)-derived hematopoietic stem and progenitor cells (HSPCs). We combined CEBPA-p30 expression with loss-of-function mutations in *GATA2*, *TET2*, or *WT1*, and evaluated the consequences of individual and combined mutations *in vitro* and *in vivo*. Through bulk and single-cell multi-omics analyses, we delineated the molecular programs underlying CEBPA-p30-driven AML and identified targetable pathways with therapeutic potential.

## Results

### CEBPA-p30 combined with loss of TET2 or WT1 leads to uncontrolled proliferation of BM HSPCs and a block in differentiation

To investigate the initiation and transformation of CEBPA-p30-driven leukemia in BM-derived CD34^+^ HSPCs from healthy young donors, we targeted the N-terminal coding region of *CEBPA* alone or in combination with the three most frequently co-mutated genes in *CEBPA*-mutant AML: *TET2*, *GATA2*, and *WT1* (Fig. 1A; Supplementary Fig. S1A) (12). For all genes, except *GATA2,* a double sgRNA strategy was used to increase editing-efficiencies. Specifically, for *WT1*, we targeted exon 7, which is a hotspot region for frame-shift mutations in AML patients (20) whereas the first exons of *TET2 and GATA2* were targeted to generate complete knockouts of these proteins. An sgRNA directed against the safe-harbor locus *AAVS1* was used as a control (21). We generally achieved indel frequencies of 90-100% for *CEBPA*, *TET2* and *WT1*. To model haploinsufficiency for *GATA2*, we titrated the sgRNA to achieve indel frequencies <50%, resulting in predominantly heterozygous loss (GATA2 knockdown (KD)) (Fig. 1B). Western blot analysis of BM CD34^+^ HSPCs 96 hours post-electroporation confirmed loss of CEBPA-p42 and concomitant upregulation of CEBPA-p30. Notably, although CEBPA-p42 loss was already evident at 24 hours, CEBPA-p30 upregulation remerged only at 96 hours, suggesting that transcriptional rewiring precedes CEBPA-p30 expression (Supplementary Fig. S1B). Loss of TET2 and WT1 protein, as well as partial loss of GATA2, was confirmed 24 hours post electroporation (Fig. 1C). The WT1 antibody recognizes the N-terminal region, therefore, exon 7 mutations lead to a complete loss of WT1, consistent with previous observations that mutated *WT1* mRNA is susceptible to nonsense-mediated RNA decay (22). Colony-forming-cell (CFC) assays revealed a reduced overall colony-forming capacity in CEBPA-p30 cells, irrespective of co-mutations and colony type, at the first plating. All CEBPA-p30 conditions, as well as TET2 KO cells, demonstrated enhanced self-renewal capacity upon serial replating (Fig. 1D). Importantly, editing efficiencies for *TET2* and *CEBPA* were 100% from first plating and remained stable across serial replating (Fig. 1E). For *WT1*, indel frequencies were 58% (WT1 KO) and 77% (WT1 KO + CEBPA-p30) at first plating and increased to 100% following the first round of CFCs. Similarly, *GATA2* indel frequencies increased from 17% (GATA2 KD) and 16% (GATA2 KD + CEBPA-p30) to ~50% efficiency after 1-2 rounds of serial replating (Fig. 1E).

**Figure 1.**
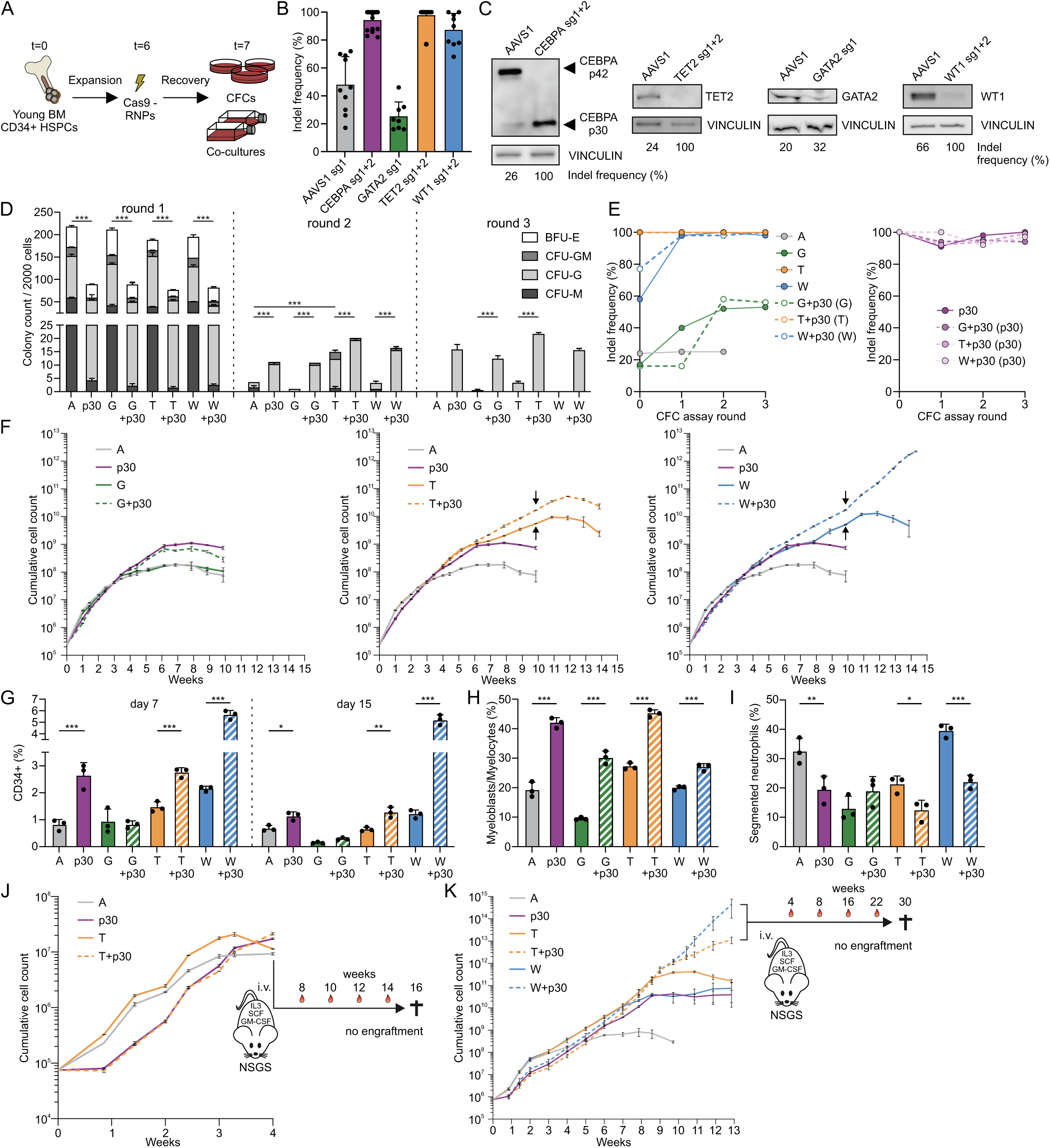
Loss of TET2 or WT1 accelerates CEBPA-p30-driven transformation. **(A)** Experimental setup of expansion and electroporation of bone marrow (BM)-derived CD34^+^ hematopoietic stem and progenitor cells (HSPCs). HSPCs were electroporated with Cas9 ribonucleoproteins (RNPs) and studied *in vitro* in colony-forming-cell (CFC) assays and co-cultures on a layer of mouse stromal-5 (MS5) cells. **(B)** Insertion/deletion (indel) frequency 24 hour (hr) after electroporation of individual BM samples. sg; sgRNA. **(C)** Western blot of CEBPA (96hr), TET2 (24hr), GATA2 (24hr), and WT1 (24hr) after electroporation. Vinculin was used as loading control. Indel frequency of the relevant gene is indicated below each lane. **(D)** Colony count of BM-derived CD34^+^ HSPCs after plating (round 1), replating (round 2), and replating twice (round 3). From hereon: A; AAVS1, p30; CEBPA-p30, G; GATA2 knockdown (KD), T; TET2 knockout (KO), and W; WT1 KO. **(E)** Indel frequency of CFCs shown in panel D. Indel frequency was determined 24hr after electroporation (t=0) and after each round of colony analysis. In case of double-mutants (dashed lines), the gene between brackets is shown. **(F)** Cumulative cell counts of edited HSPCs grown on a stromal layer of MS5 cells. Arrows indicate when cells were replated onto a fresh layer of MS5 cells. **(G)** Percentage of CD34^+^ cells in the supernatant harvested at day 7 (left) and day 15 (right). **(H-I)** Percentage of myeloblasts/myelocytes **(H)** and segmented neutrophils **(I)** in the supernatant harvested at day 15. **(J-K)** Cumulative cell counts of two independent HSPC co-cultures. Supernatant and adherent cells of A, C, T, and C+T **(J)** or C+T and C+W **(K)** cultures were harvested after 4 weeks or 13 weeks respectively and transplanted intravenously (i.v.) into NOD.Cg-Prkdc^scid^ Il2rg^tm1Wjl^ Tg(CMV-IL3,CSF2,KITLG)1Eav/MloySzJ (NSGS) mice. Bar plots represent mean ± SD. Error bars in growth curves represents mean ±SD of technical triplicates. *p<0.05, **p<0.01, ***p<0.001.

To assess proliferation dynamics and lineage output over time, we co-cultured gene-edited HSPCs in Gartner’s medium supplemented with thrombopoietin (TPO), stem cell factor (SCF), interleukin-3 (IL-3), and granulocyte colony-stimulating factor (G-CSF) on an adherent layer of mouse stromal-5 (MS5) cells (23). Consistent with findings from the CFC assays, indel frequencies for *CEBPA*, *TET2*, and *WT1* were already high at plating and rapidly reached 100%, whereas *GATA2* indel frequencies increased to ~50% after 8 weeks in culture (Supplementary Fig. S1C). At early timepoints, CEBPA-p30 cells displayed reduced proliferation and elevated frequencies of CD34^+^ HSPCs regardless of co-mutational status (Fig. 1F-G; Supplementary Fig. S1D). AAVS1 control and GATA2 KD cells showed decreased proliferation capacity after 4-5 weeks, whereas CEBPA-p30, TET2 KO, and WT1 KO cells maintained significantly prolonged proliferation. Strikingly, combining TET2 KO or WT1 KO with CEBPA-p30 drove uncontrolled proliferation, sustained up to 12 and 15 weeks, respectively (Fig. 1F). These cells were phenotypically immature, with increased proportions of myeloblasts/myelocytes and reduced mature segmented neutrophils (Fig. 1H-I; Supplementary Fig. S1E). When co-cultured cells were transferred to liquid culture after 10 weeks, only WT1 KO + CEBPA-p30 continued to expand without stromal support, exhibiting a complete differentiation block at the myeloblastic stage (Supplementary Fig. S1F-H). Together, these data indicate that CEBPA-p30 in combination with TET2 or WT1 loss can drive leukemic transformation, characterized by uncontrolled proliferation and impaired granulocytic differentiation. To test whether these transformed cells could establish AML *in vivo*, we transplanted the combined adherent and suspension fractions from two independent co-cultures derived from two independent BM donors into NOD.Cg-Prkdc^scid^ Il2rg^tm1Wjl^ Tg(CMV-IL3,CSF2,KITLG)1Eav/MloySzJ (NSGS) mice, either after 4 or 13 weeks of culture (Fig. 1J-K). Despite resembling leukemic transformation *in vitro*, none of these conditions led to detectable human (hCD45^+^) engraftment (data not shown). This may reflect insufficient functional leukemic stem cells (LSCs), resulting from loss of LSC activity during prolonged culture and the intrinsically low frequency of engraftable LSCs in CEBPA-mutant AML (24).

### CEBPA-p30 cells depend on human cytokines and the BM microenvironment

Since AML could not be established from long-term *in vitro* cultured cells, we transplanted BM HSPCs directly after gene editing into NOD.Cg-Prkdc^scid^ Il2rg^tm1Wjl^ (NSG) mice (Fig. 2A). AAVS1 cells engrafted consistently, maintaining an average of 10% peripheral blood (PB) chimerism and stable indel frequency over 30 weeks (Fig. 2B-C). In this experiment, we used only one sgRNA against *TET2*, resulting in modest editing efficiencies at injection. Nonetheless, TET2 KO cells expanded significantly in both the single- and double-mutant context, with increased PB engraftment upon TET2 loss, as also observed previously in human TET2-deficient cells (25). In contrast, CEBPA-p30 and TET2 KO + CEBPA-p30 cells showed markedly reduced PB engraftment compared to AAVS1 and TET2 KO cells (Fig. 2B-C). The few CEBPA-p30 cells detectable in PB exhibited a myeloid bias at the expense of lymphoid differentiation (Fig. 2D-E). At sacrifice, BM hCD45^+^ engraftment was comparable between AAVS1 and CEBPA-p30 mice, whereas *TET2* single-mutant mice showed increased BM engraftment (Fig. 2F-G). Splenic engraftment of CEBPA-p30 cells was negligible, consistent with poor PB dissemination (Fig. 2H). BM-derived CEBPA-p30 cells showed HSPC frequencies comparable to AAVS1 controls but exhibited a myeloid bias and reduced lymphoid differentiation (Fig. 2I-L). These findings were confirmed in an independent BM donor experiment that also included WT1 KO and GATA2 KD conditions (Supplementary Fig. S2A-F). Interestingly, WT1 KO cells showed increased BM and PB engraftment, similar to TET2 KO cells (Supplementary Fig. S2A-B). In contrast, high *GATA2* indel frequency after electroporation diminished engraftment of *GATA2* single- and double-mutant cells, consistent with homozygous *Gata2* loss causing HSC exhaustion (Supplementary Fig. S2A-C) (26).

**Figure 2.**
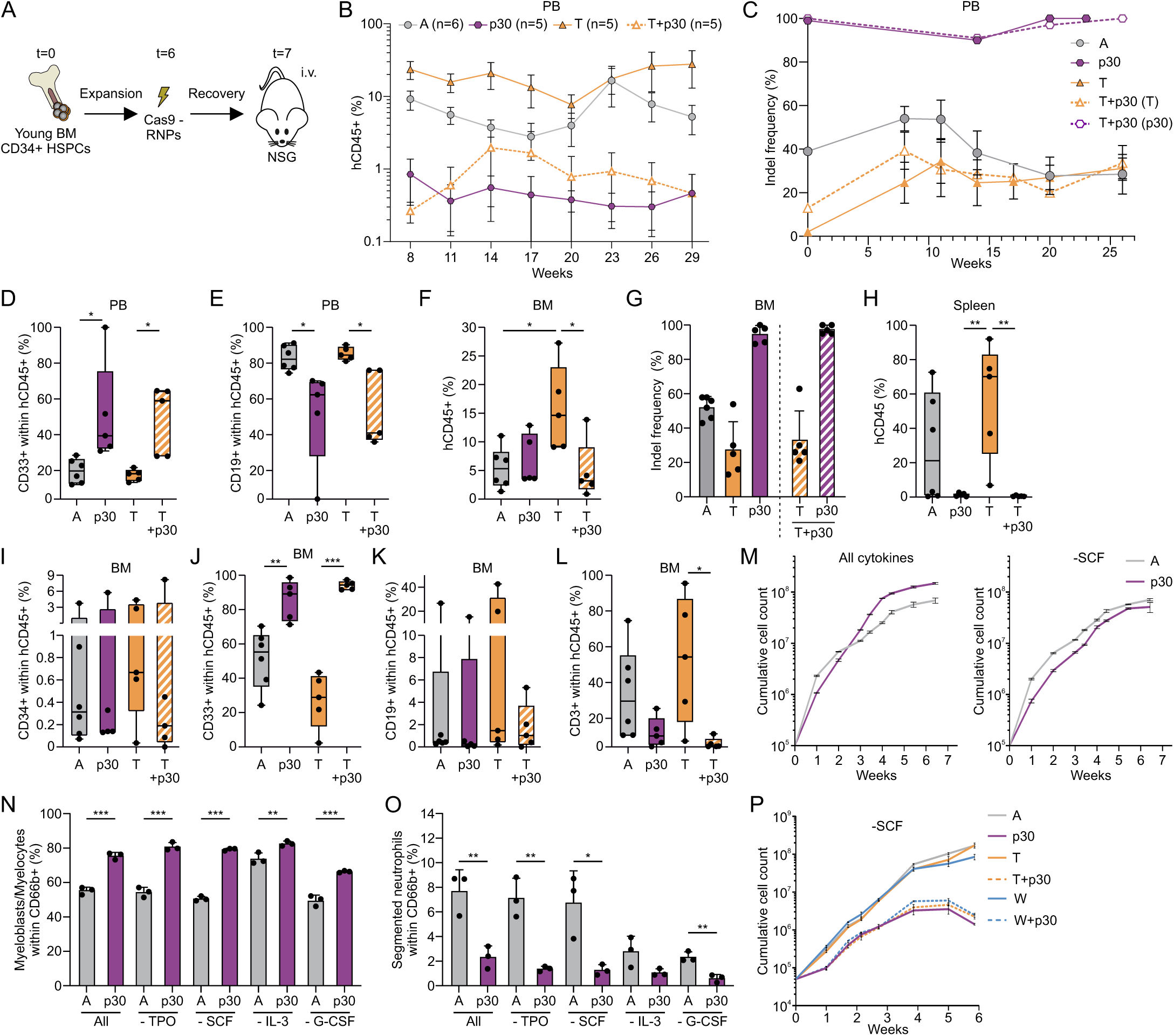
CEBPA-p30 cells rely on human cytokines and interaction with the BM microenvironment. **(A)** Experimental setup of BM-derived CD34^+^ HSPC expansion, subsequent electroporation with RNPs, and i.v. transplantation into sub-lethally irradiated NOD.Cg-Prkdc^scid^ Il2rg^tm1Wjl^ (NSG) mice. **(B-C)** Engraftment of human CD45^+^ (hCD45^+^) cells **(B)** and indel frequency within hCD45^+^ cells **(C)** in peripheral blood (PB) of NSG mice through time. **(D-E)** Percentage of CD33^+^ **(D)** and CD19^+^ **(E)** cells within hCD45^+^ cells in the PB at week 8. **(F)** Percentage of hCD45^+^ cells in the BM at sacrifice. **(G)** Indel frequency in hCD45^+^CD33^+^ cells isolated from the BM of NSG mice at sacrifice. **(H)** Percentage of hCD45^+^ cells in the spleen at sacrifice. **(I-L)** Percentage of CD34^+^ **(I)**, CD33^+^ **(J)**, CD19^+^ **(K)**, and CD3^+^ **(L)** cells within BM-derived hCD45^+^ cells harvested at sacrifice. **(M)** Cumulative cell counts of edited HSPCs grown on a stromal layer of MS5 cells in medium containing thrombopoietin (TPO), stem cell factor (SCF), interleukin-3 (IL-3), and granulocyte-colony stimulating factor (G-CSF) (All, left) or lacking SCF (-SCF, right). **(N-O)** Percentage of myeloblasts/myelocytes **(N)** and segmented neutrophils **(O)** in the supernatant at week 4 in the presence of all cytokines or lacking one cytokine. **(P)** Cumulative cell counts of edited HSPCs grown on a stromal layer of MS5 cells in the presence of TPO, IL-3, and G-CSF but lacking SCF. Boxplots show median (line), interquartile range (box), and minimum/maximum values (whiskers). Bar plots represent mean ± SD. Error bars in growth curves represents mean ± SD of technical triplicates. *p<0.05, **p<0.01, ***p<0.001.

Murine *Cebpa^N/N^* cells have been reported to depend on SCF and IL-3 for proliferation (8). Since CEBPA-p30 cells expanded robustly in our MS5 co-culture system (containing SCF, TPO, IL-3, and G-CSF) but not in NSG mice (which lack human cytokines), we used the co-culture system to determine whether this proliferative advantage was cytokine-dependent. CEBPA-p30 cells required SCF and G-CSF for their proliferative advantage, whereas IL-3 and TPO played lesser roles (Fig. 2M; Supplementary Fig. S2G). Across all cytokine conditions, CEBPA-p30 cells maintained high indel frequency and a differentiation block marked by increased myeloblasts/myelocytes and reduced segmented neutrophils (Fig. 2N-O; Supplementary Fig. S2L). Loss of TET2 or WT1 in combination with CEBPA-p30 did not rescue the phenotype observed in the absence of SCF (Fig. 2P). In summary, while CEBPA-p30 cells engraft in the BM of NSG mice and display a myeloid bias, they fail to disseminate systemically and depend on SCF and G-CSF for sustained proliferation.

### CEBPA-p30 combined with TET2 or WT1 loss drives AML development

Previous work has demonstrated that primary AML samples engraft more efficiently in NSGS mice, which express human SCF, IL-3, and GM-CSF, compared to NSG mice (27). Given the observed dependency of CEBPA-p30 cells on SCF and G-CSF, we tested whether AML could be established *in vivo* by transplanting freshly gene-edited BM HSPCs into NSGS mice (Fig. 3A). In contrast to their poor PB dissemination in NSG mice, CEBPA-p30 cells showed PB chimerism similar to AAVS1 controls in NSGS mice and retained their myeloid bias (Fig. 3B-C). Before the 12-week post-transplant experimental endpoint, two mice of the TET2 KO + CEBPA-p30 condition and one mouse of the WT1 KO + CEBPA-p30 condition developed lethal AML (Fig. 3D). The remaining mice of these groups exhibited clear signs of AML at sacrifice, including pale bones, high hCD45^+^ engraftment in BM and spleen, and splenomegaly (Fig. 3E-F; Supplementary Fig. S3A-B). Editing efficiencies for *CEBPA*, *TET2*, and *WT1* reached 100% by the time of sacrifice, while *GATA2* editing was established at ~10% and increased to ~30% (Fig. 3G). Phenotypic characterization confirmed increased myeloblasts/myelocytes and monocytes in CEBPA-p30 samples, along with reduced megakaryocytic and B-cell output, while HSPC frequencies remained comparable (Fig. 3H-M). Secondary transplantation of leukemic cells into NSGS mice failed to engraft, likely due to HSC exhaustion under high cytokine exposure (data not shown) (28). Incubation longer than 12 weeks post-transplantation resulted in macrophage activation syndrome (MAS) caused by emerging T-cell subsets, especially in the non-CEBPA-p30 conditions (data not shown) (29). To minimize confounding effects caused by MAS, two subsequent NSGS experiments were terminated 12 weeks post-transplantation. Importantly, AML formation from cells expressing CEBPA-p30 combined with loss of TET2 or WT1 was very reproducible with two independent BM donors across two mouse cohorts, despite variability in donor engraftment capacity as seen from the AAVS1 controls (Supplementary Fig. S3C-J). Together, these findings establish a robust *in vivo* model of *CEBPA*-mutant AML using CRISPR-edited primary young BM HSPCs. This platform enables investigation of human CEBPA-p30 biology and the role of common co-mutated genes in *CEBPA*-mutant leukemias.

**Figure 3.**
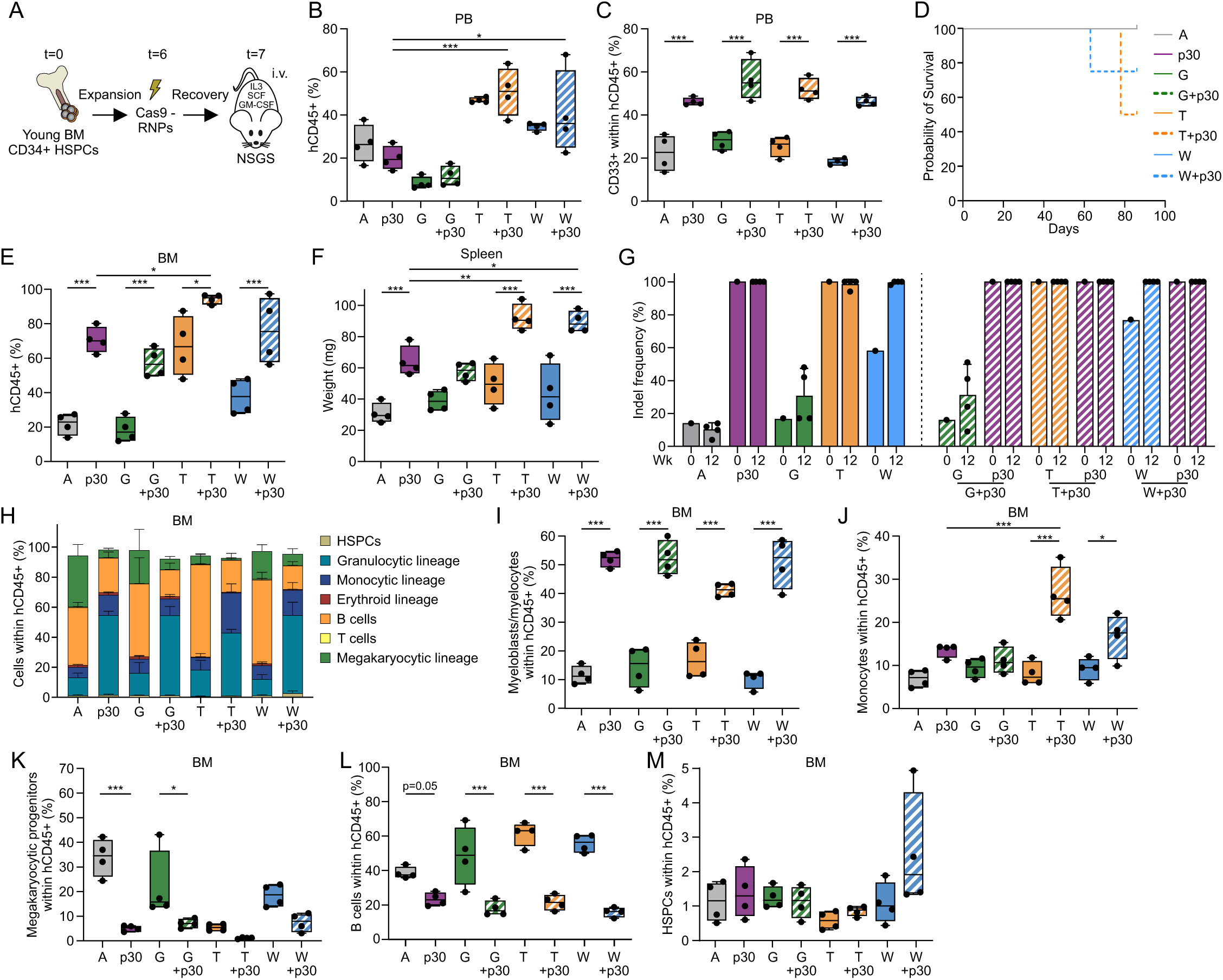
CEBPA-p30 combined with TET2 or WT1 loss induces lethal AML in NSGS mice. **(A)** Experimental setup of BM-derived CD34^+^ HSPC expansion, subsequent electroporation with RNPs, and i.v. transplantation into sub-lethally irradiated NSGS mice. **(B-C)** Percentage of hCD45^+^ cells **(B)** and CD33^+^ cells within the hCD45^+^ compartment **(C)** in PB at week 8. **(D)** Kaplan-Meier curve of mice transplanted with cells from eight different conditions. **(E)** Percentage of hCD45^+^ cells in the BM at sacrifice. **(F)** Spleen weight at sacrifice. **(G)** Indel frequency at time of transplantation (week (wk) 0) and in BM hCD45^+^CD3^−^ CD19^−^ cells at sacrifice (wk 12). T+p30 and W+p30 mice that were sacrificed before the experimental endpoint are included at the wk 12 timepoint. **(H)** Distribution of different hematopoietic lineages in BM-derived hCD45^+^ cells at sacrifice. **(I-M)** Percentage of myeloblasts/myelocytes (CD66b^+^CD16^−^) **(I)**, monocytes (CD14^+^CD11b^+^) **(J)**, megakaryocytic progenitors (CD41a^+^CD117^+^) **(K)**, B cells (CD19^+^) **(L)**, and HSPCs (CD34^+^CD3^−^CD19^−^) **(M)** within BM-derived hCD45^+^ cells at sacrifice. Boxplots show median (line), interquartile range (box), and minimum/maximum values (whiskers). *p<0.05, **p<0.01, ***p<0.001.

### CEBPA-p30, but not CEBPA^C/C^ mutations drives the myeloid bias

Recent studies demonstrate that patients with *CEBPA^C^* mutations have a favorable prognosis, irrespective of co-occurring *CEBPA^N^* mutations, and these patients form a distinct subgroup in the most recent European LeukemiaNet classification (5,30–32). To test whether the observed CEBPA-p30 phenotypes could be replicated by disruption of the CEBPA bZIP domain, we designed two sgRNAs targeting the bZIP domain of *CEBPA* (CEBPA^C/C^), mimicking a small in-frame deletion as also observed in patients. The resulting cells were transplanted into NSGS mice alongside mice transplanted with AAVS1 or CEBPA-p30 cells (Fig. 4A-B). Whereas the CEBPA-p30 cells again showed significantly increased PB chimerism compared to AAVS1 controls, CEBPA^C/C^ cells exhibited reduced chimerism (Fig. 4C). Both CEBPA-p30 and CEBPA^C/C^ cells displayed a myeloid bias, as indicated by the elevated CD33 expression (Fig. 4D). At sacrifice, CEBPA-p30 cells again demonstrated enhanced BM engraftment, whereas CEBPA^C/C^ mice were comparable to AAVS1 controls (Fig. 4E). *AAVS1* editing efficiencies remained stable over time while CEBPA-p30 editing reached 100% at sacrifice. *CEBPA^c/c^*gene editing produced a mixture of in-frame and out-of-frame deletions with a modest increase in edited cells at sacrifice, but no specific clonal pattern emerged and neither mutation type conferred a selective engraftment advantage (Fig. 4F-G). Phenotypic characterization revealed that CEBPA-p30 cells had increased myeloblasts/myelocytes and monocytes, and reduced megakaryocytic cells, consistent with previous results (Fig. 4H-L). In contrast, CEBPA^C/C^ cells exclusively generated megakaryocytic progenitors co-expressing CD33, CD41a, and CD117, suggesting that loss of functional CEBPA-p42 without accompanying CEBPA-p30 expression restricts differentiation to the megakaryocytic lineage in NSGS mice, in line with findings from murine *Cebpa^C^* models (Fig. 4H-L) (9). Similar results were observed in spleen (Fig. 4M-O). Overall, these findings suggest that functional loss of CEBPA-p42 without concomitant CEBPA-p30 expression is insufficient to drive granulocytic/monocytic commitment or initiate myeloid leukemia.

**Figure 4.**
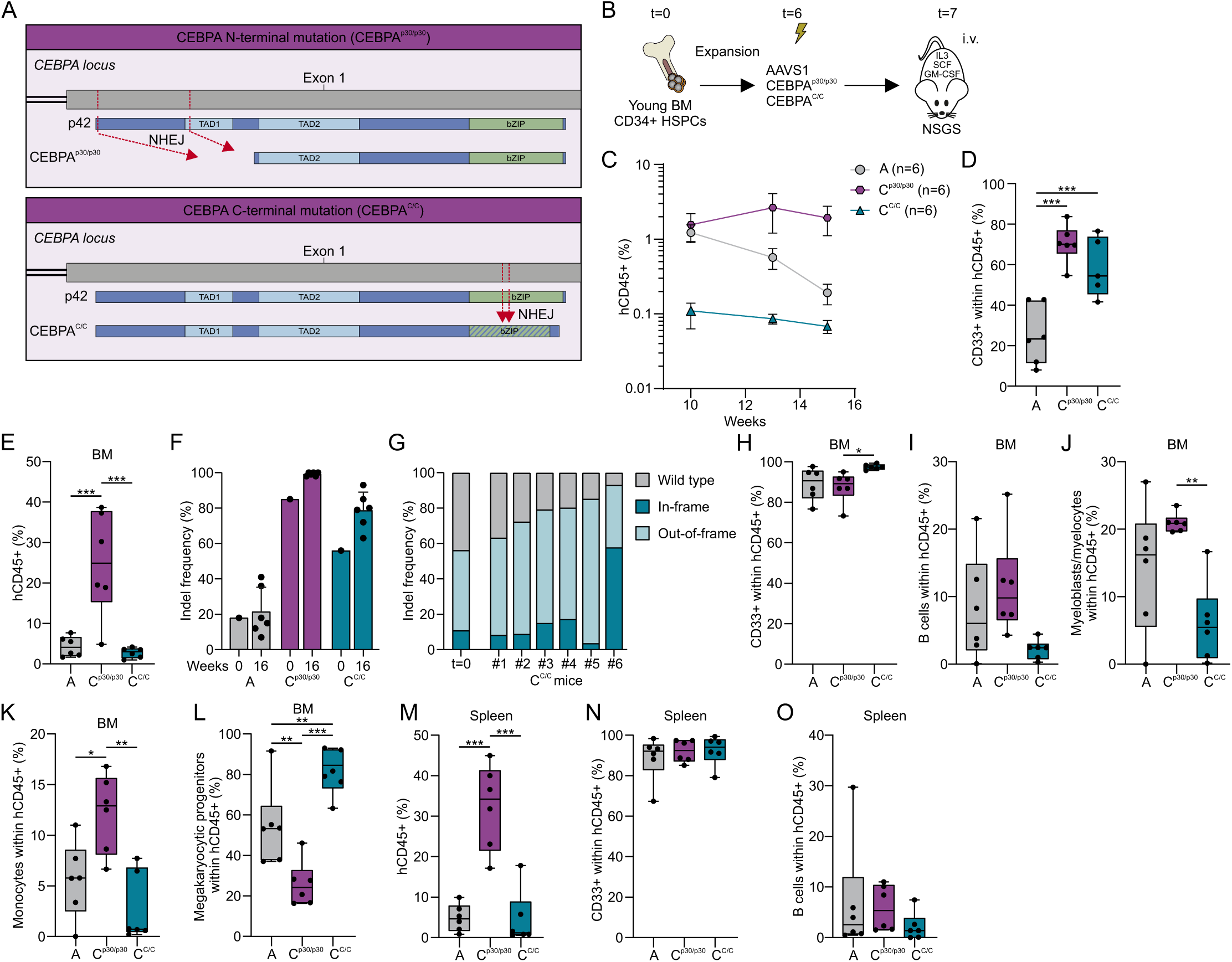
CEBPA-p30, but not CEBPA^C/C^, induces monocytic/granulocytic bias. **(A)** Schematic representation of sgRNA binding sites within the N-terminal and C-terminal coding regions of CEBPA in CEBPA-p30 (top) and CEBPA^C/C^ (bottom) models. bZIP; Basic Leucine Zipper, NHEJ; non-homologous enjoining, TAD; transactivation domain. **(B)** Experimental setup of BM-derived CD34^+^ HSPC expansion, subsequent electroporation with RNPs, and i.v. transplantation into sub-lethally irradiated NSGS mice. **(C)** Percentage of hCD45^+^ cells in the PB measured over time. **(D)** Percentage of CD33^+^ cells within PB hCD45^+^ at week 10. **(E)** Engraftment of hCD45^+^ cells in the BM at sacrifice. **(F)** Indel frequency at day of transplantation (wk 0) and at sacrifice (wk 16) in BM hCD45^+^CD3^−^CD19^−^ cells. **(G)** Percentage of wild type, in-frame, and out-of-frame alleles at the C-terminal coding region of *CEBPA* in the injected cells at time of injection (t=0) and at sacrifice in individual CEBPA^C/C^ mice. **(H-L)** Percentage of CD33^+^ cells **(H)**, B cells (CD19^+^) **(I)**, myeloblasts/myelocytes (CD66b^+^CD16^−^) **(J)**, monocytes (CD14^+^CD11b^+^) **(K)**, and megakaryocytic progenitors (CD41a^+^CD117^+^) **(L)** within BM hCD45^+^ cells at sacrifice. **(M)** Percentage of hCD45^+^ cells in the spleen at sacrifice. **(N-O)** Percentage of CD33^+^ cells **(N)** and B cells (CD19^+^) **(O)** within hCD45^+^ cells in the spleen at sacrifice. Boxplots show median (line), interquartile range (box), and minimum/maximum values (whiskers). *p<0.05, **p<0.01, ***p<0.001.

### CEBPA-p30 drives a myeloid bias and expansion of a conventional dendritic type 2 (cDC2) like subset

To further investigate the mechanism behind the myeloid bias caused by CEBPA-p30, we performed single-cell RNA sequencing (scRNA-seq) using an on-chip multiplexing approach. For each condition, equal ratios of hCD45^+^CD3^−^CD19^−^ and hCD45^+^CD3^−^CD19^−^CD34^+^ cells were sorted from two individual NSGS mice (Fig. 5A; Supplementary Fig. S4A). Indel frequencies within the CD34^+^ fraction were close to 100% for most conditions and ~40% for *GATA2*, comparable with indel frequencies in total cell fractions (Fig. 3G; Supplementary Fig. S4B). Using condition-specific barcodes, ~4000 cells could be assigned per condition, with on average ~5000 protein coding genes detected per cell (Supplementary Fig. S5A-B, Supplementary Data 1). To gain a first impression of leukemic cell identity, we mapped CEBPA-p30 double-mutant conditions onto a single-cell reference atlas of healthy hematopoiesis (Supplementary Fig. S5C) (33). CEBPA-p30 cells in combination with TET2 KO, WT1 KO, or GATA2 KD aligned with conventional dendritic cells (cDCs) and monocytic lineages, consistent with scRNA-seq profiles of primary *CEBPA*-mutant leukemias (33). We then generated a reference map by projecting all conditions onto one UMAP and annotating clusters based on lineage-specific gene expression (Fig. 5B; Supplementary Fig. S5D-E) (34–39). Myeloid clusters expressed both monocytic and granulocytic genes, including *AZU1*, *CD14*, and *CEACAM8*, without clear separation, likely reflecting limitations of immunocompromised mouse models in supporting full granulocytic maturation and technical challenges in capturing mature granulocytes by scRNA-seq (40). On the integrated map, CEBPA-p30 cells were enriched along the monocytic/granulocytic lineage at the expense of basophil/eosinophil/mast cell (BaEoMa) precursors. This myeloid skew was further enhanced in combination with *TET2*, *WT1*, or *GATA2* mutations, consistent with flow cytometry observations (Fig. 5C-D). In addition, CEBPA-p30 conditions exhibited a lineage shift from cDC type 3 (cDC3) toward cDC2-like cells, characterized by expression of cDC-associated *CDC1* and AML-associated genes including *CD117* and *IL1RAP* (Fig. 5C-D; Supplementary Fig. S5E-F) (41–43). Notably, GATA2 KD + CEBPA-p30 cells also showed a marked increase in pro-erythroblasts alongside the myeloid bias, highlighting a potential source of cells capable of initiating acute erythroid leukemia (AEL), as observed in patients and mouse models with this specific mutational combination (Fig. 5C-D; Supplementary Fig. S5E) (13,44). Despite similar proportions of early HSPCs, *CEBPA* expression was upregulated in the HSC/MPP cluster across all CEBPA-p30 conditions, suggesting that lineage fate decisions in *CEBPA*-mutant cells are established at an early stage (Fig. 5E).

**Figure 5.**
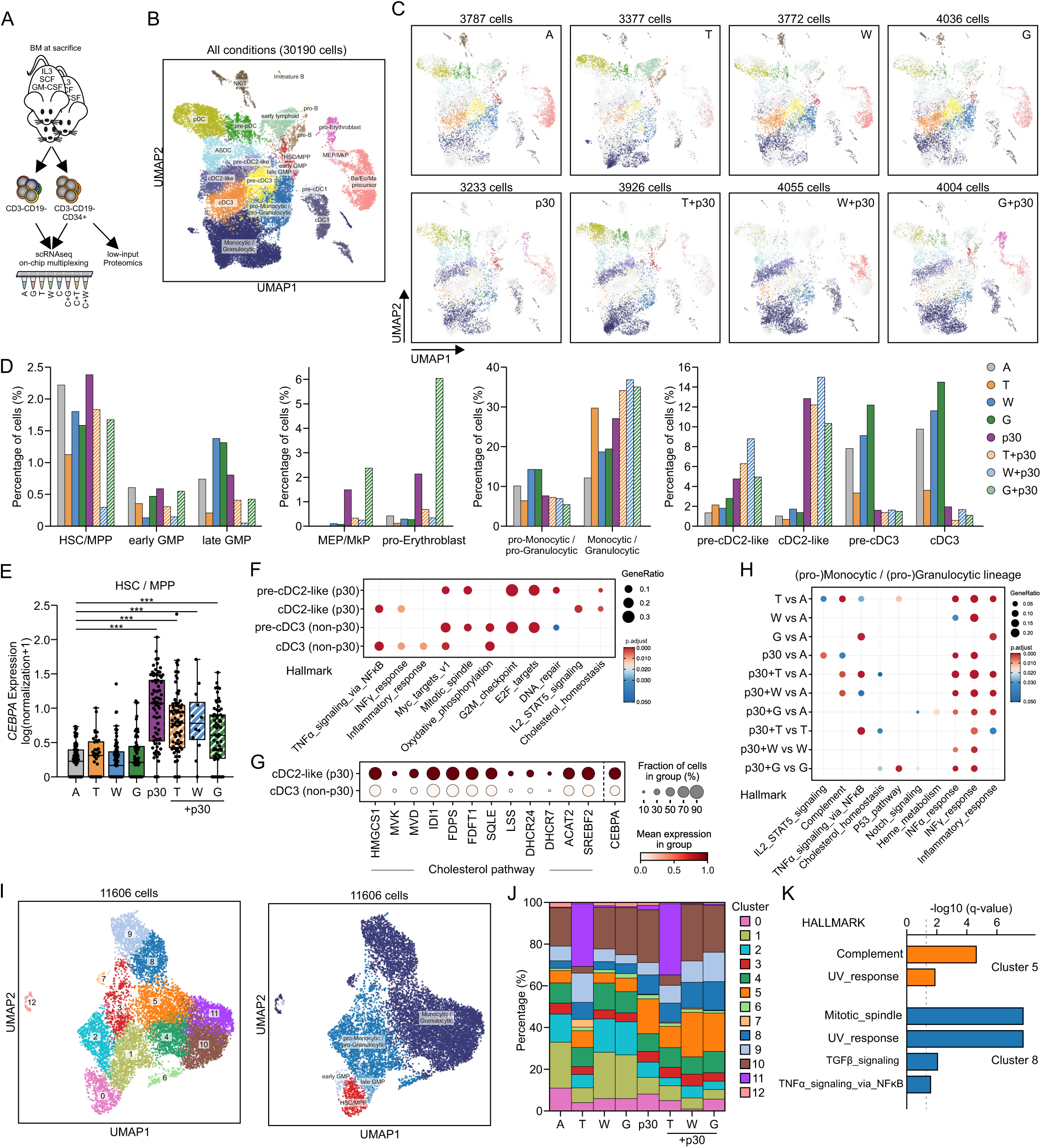
scRNA-seq confirms CEBPA-p30-driven myeloid bias and reveals a shift toward cDC2-like cells. **(A)** scRNA-seq with on-chip multiplexing was performed on equal ratios of BM-derived CD34^+^CD3^−^CD19^−^ HSPCs and hCD45^+^CD3^−^CD19^−^ from two mice of each condition. **(B)** UMAP of all conditions with cluster annotations. **(C)** UMAPs showing individual conditions overlaid on the combined UMAP from (B). **(D)** Percentage of given cell types including immature HSPCs and myelo-erythroid lineages within each condition. **(E)** *CEBPA* expression within the HSC/Multipotent Progenitor (MPP) cluster. **(F)** Pathway enrichment analysis of CEBPA-p30 (pre-)cDC2-like and non-CEBPA-p30 (pre-)cDC3 cells. **(G)** Expression of cholesterol-associated genes in CEBPA-p30 cDC2-like and non-CEBPA-p30 cDC3 cells. **(H)** Pathway enrichment analysis of CEBPA-p30-enriched pathways in pooled (pro-)monocytic/(pro-)granulocytic clusters. **(I)** Re-clustered UMAP containing only the HSC/MPP, early and late granulocyte-monocyte progenitors (GMP), and (pro-)monocytic/(pro-)granulocytic clusters as defined in (B). **(J)** Cell type distribution within each condition using clusters from (I). **(K)** Pathway enrichment analysis of CEBPA-p30-enriched pathways in clusters 5 and cluster 8. ***p<0.001.

### CEBPA-p30-driven AML shows increased IFN signaling and cholesterol biosynthesis

To explore pathways driving the CEBPA-p30 phenotype, we performed pathway enrichment analysis on the (pre-)cDC2-like and (pre-)cDC3 clusters. While both were enriched for inflammatory signatures, IL2-STAT5 signaling and cholesterol homeostasis were specifically enriched in cDC2-like cells compared to all other cell types (Fig. 5F). Consistently, cDC2-like cells expressed high levels of genes directly involved in the pathway converting acetyl-CoA to cholesterol (Fig. 5G). Enrichment analysis on the monocytic/granulocytic compartment further revealed upregulation of interferon (IFN)-α and IFN-γ response pathways across all CEBPA-p30 conditions (Fig. 5H). TET2 KO cells exhibited a strong IFN signature independent of CEBPA-p30, whereas GATA2 KD + CEBPA-p30 cells were uniquely enriched for heme metabolism. In addition, both TET2 KO + CEBPA-p30 and GATA2 KD + CEBPA-p30 cells displayed enrichment for cholesterol homeostasis pathways (Fig. 5H). To examine the myelocytic-granulocytic populations in greater detail, we re-clustered this compartment, which accounted for one-third of the full reference map (Fig. 5I). Marker gene expression delineated a trajectory from immature (cluster 0) to more mature states (clusters 8-11) (Fig 5I and Supplementary Fig. S6A). Clusters 5 and 8, enriched in CEBPA-p30 conditions, were characterized by tumor necrosis factor alpha (TNF-α), transforming growth factor beta (TGF-β), and complement pathway signatures (Fig. 5J-K; Supplementary Fig. S6B). Cluster 11 was unique to *TET2* single- and double-mutants and was strongly enriched for IFN-α/γ response genes and high *CEACAM8* expression, a marker specific to granulocytic cells (Fig. 5J; Supplementary Fig. S6A-C). This aligns with recent findings describing a highly inflamed granulocytic subset of TET2 KO cells (45). Together, these data show that CEBPA-p30 promotes cDC2-like differentiation at the expense of cDC3 cells and myeloid lineages characterized by inflammatory and cholesterol-related transcriptional programs. Partial loss of GATA2 in a CEBPA-p30 context additionally enhances erythroid potential, while TET2 loss promotes expansion of an inflamed granulocytic subset with strong interferon signatures.

To further substantiate these findings, we performed low-input proteomics on only 500 hCD45^+^CD3^−^CD19^−^CD34^+^ HSPCs from three mice per condition (Fig. 5A, Supplemental Figure S4A-B). Across samples, we annotated ~6500 proteins (up to four technical replicates averaged per mouse) (Supplementary Fig. 7A and Supplementary Data 2). Principal component analysis revealed that CEBPA-p30 and loss of TET2, GATA2, or WT1 each induced prominent proteomic changes, with double-mutant conditions exhibiting unique proteomic profiles distinct from both AAVS1 controls and their respective single-mutants (Fig. 6A-C). TET2 loss activated IFN signaling, WT1 loss affected mitochondrial processes, and partial GATA2 loss altered amino-acid metabolism (Supplementary Fig. 7D-O, Supplementary Data 2). Focusing on proteins upregulated in CEBPA-p30 samples relative to AAVS1 controls, we observed consistent enrichment of neutrophil differentiation and cholesterol biosynthesis pathways across all CEBPA-p30 conditions, regardless of co-mutations (Fig. 6D-E, Supplementary Data 2). No clear overlapping processes were observed among the proteins downregulated in CEBPA-p30 cells (Supplementary Fig. 7B-C and Supplementary Data 2). In the context of CEBPA-p30, loss of TET2 or WT1 further enhanced IFN signaling and increased expression of key proteins associated with major histocompatibility complex (MHC) class I and II biology. By contrast, combining CEBPA-p30 with partial GATA2 loss induced a unique program enriched for proteins involved in ribosomal and messenger RNA processing (Fig. 6D-E). Consistent with our scRNAseq findings, many enzymes directly involved in cholesterol biosynthesis from acetyl-CoA were upregulated across all CEBPA-p30 conditions, with the strongest induction observed upon *WT1* co-mutation (Figure 6F).

**Figure 6.**
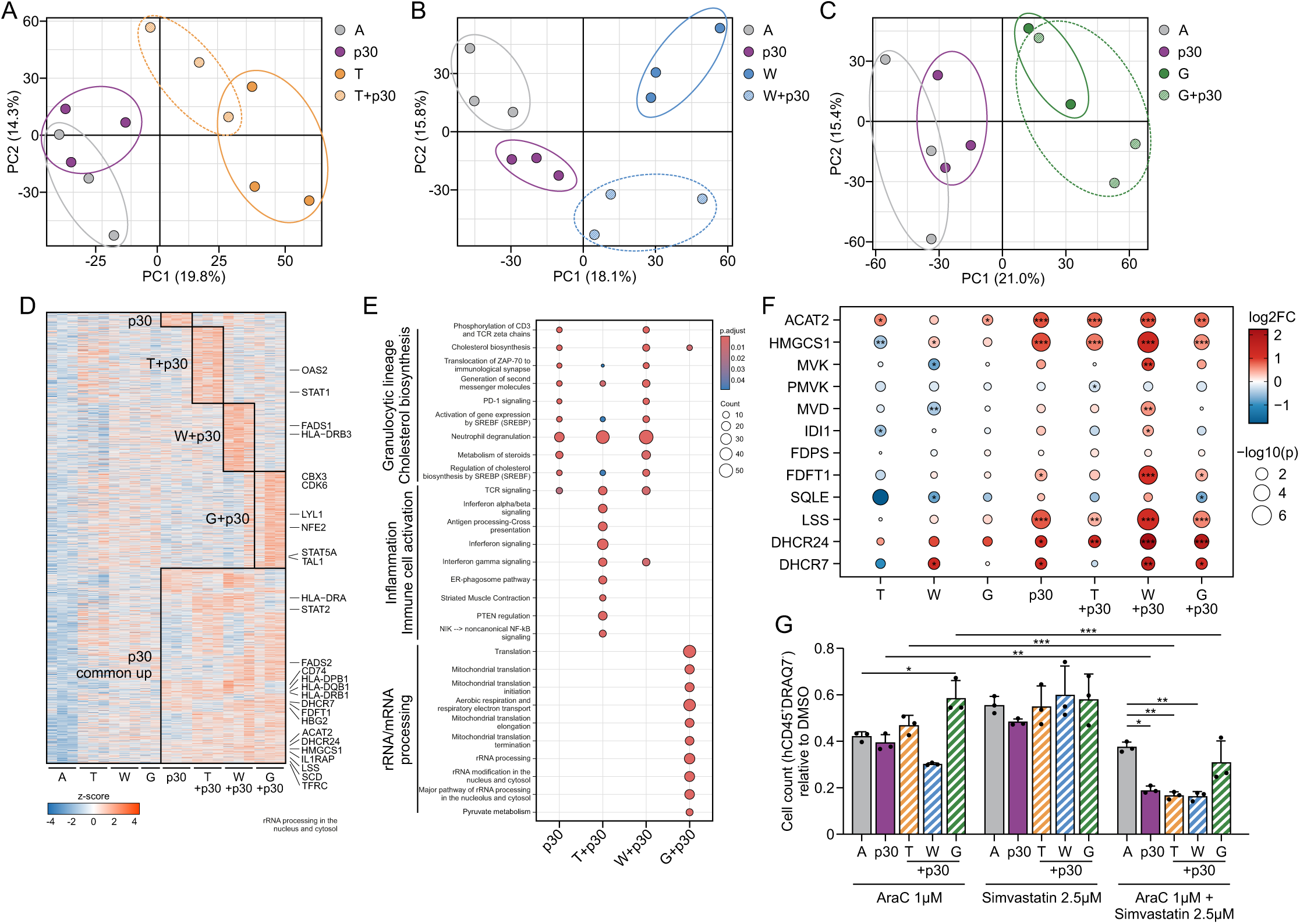
CEBPA-p30 HSPCs show elevated cholesterol biosynthesis protein levels. **(A-C)** Principal component analysis of protein abundance in BM-derived hCD45^+^CD34^+^CD3^−^CD19^−^ HSPCs of A, T, p30, and T+p30 **(A)**, A, W, p30, and W+p30 **(B)**, and A, G, p30, and G+p30 **(C)**. **(D)** Proteins upregulated (log2FC > log2(1.3), p < 0.05) in CEBPA-p30 single- and/or double-mutants relative to AAVS1 HSPCs. **(E)** Reactome pathways enriched among proteins upregulated in CEBPA-p30 single- and/or double-mutants relative to AAVS1 HSPCs. **(F)** Expression of cholesterol biosynthesis-related proteins in all conditions relative to AAVS1 HSPCs. **(G)** Supernatant cells harvested at day 30 from gene-edited BM CD34^+^ MS5 co-cultures were treated with cytarabine (AraC), simvastatin, or both for 96 hr. Bar plot shows total viable (hCD45^+^DRAQ7^−^) cell counts relative to DMSO control (set to 1). Error bars indicate mean ± SD of technical triplicates. *p<0.05, **p<0.01, ***p<0.001.

### Targeting cholesterol biosynthesis increases chemosensitivity of CEBPA-p30 AML

Increased cholesterol biosynthesis has been suggested to reduce chemosensitivity (46). Given the upregulation of this pathway across all CEBPA-p30 mutants, we tested whether inhibiting cholesterol biosynthesis could enhance chemotherapy response. We treated cells with simvastatin, an HMG-CoA reductase inhibitor that blocks the rate-limiting step converting HMG-CoA to mevalonate, in combination with cytarabine. Primary cells harvested from mice did not expand *ex vivo* (data not shown), therefore, we used our established MS5 co-culture system to generate viable cells for treatment. As previously observed (Fig. 1), co-cultured HSPCs expanded robustly, and supernatant cells proliferated in liquid culture for up to 8 days (Supplementary Fig. 8A-D). Treatment with either cytarabine or simvastatin for 96 hours reduced viable cell numbers across all conditions in two independent BM donors, with CEBPA-p30 combined with TET2 or WT1 loss showing the highest sensitivity (Fig. 6G, Supplementary Fig. 8E). However, whereas combining the two drugs did not further decrease viable cell numbers in AAVS1 control cells, all CEBPA-p30 cells showed a further reduction relative to controls (Fig. 6G, Supplementary Fig. 8E). These data suggest that *CEBPA*-mutant AML patients, especially those with *TET2* or *WT1* mutations who show reduced chemotherapy response, may benefit from cholesterol biosynthesis inhibition to enhance chemosensitivity.

## Discussion

Precise gene editing of healthy primary BM-derived HSPCs enabled transformation into overt *CEBPA*-mutant AML. Expression of CEBPA-p30 alone induced a myeloproliferative phenotype, which progressed to AML when combined with *TET2* or *WT1* mutations. The aberrant myeloid differentiation, including the shift from cDC3 cells towards cDC2-like cells and the increase in cholesterol biosynthesis, was primarily driven by CEBPA-p30. *TET2* and *WT1* mutations further activated IFN signaling pathways and enhanced CEBPA-p30-driven phenotypes, whereas partial GATA2 loss in the context of CEBPA-p30 predominantly affected rRNA and mRNA processing, resulting in increased erythroid precursors. Finally, we show that inhibiting cholesterol biosynthesis with simvastatin enhances chemosensitivity of transformed CEBPA-p30 cells.

Targeting cholesterol homeostasis has yielded promising results in AML clinical trials, yet the upstream drivers of elevated cholesterol remain poorly defined (46). A recent study demonstrated that CEBPA regulates fatty-acid metabolism and cholesterol homeostasis, including transcriptional activation of *DHCR7* and *DHCR24* downstream of the Fms-like tyrosine kinase 3 (FLT3) receptor in *FLT3*-mutant AML cells (47). Consistent with this, *FLT3-*mutant AML cells show increased sensitivity to simvastatin (48). Given the substantial overlap between CEBPA-p42 and p30 DNA-binding sites, it is likely that CEBPA-p30 regulates cholesterol biosynthesis in a manner similar to CEBPA-p42 (11,49). Importantly, we see elevated *CEBPA* expression in HSPCs of all CEBPA-p30 conditions potentially explaining the enhanced cholesterol biosynthesis. Both cholesterol biosynthesis and inflammatory signaling were further enhanced upon TET2 or WT1 loss, potentially contributing to leukemic transformation. Although IFN signaling can reduce cholesterol biosynthesis by interfering with the SCAP/SREBP2 axis, this negative feedback appears overridden in the CEBPA-p30 double-mutant cells, which maintain high cholesterol biosynthesis despite strong inflammatory cues (50). Deficient AP-1 activation in CEBPA-p30 cells may alter cellular stress responses to inflammation and contribute to sustained cholesterol production (17).

Recent cohort studies indicate that only *CEBPA^C^* variants are associated with a favorable prognosis, suggesting that these lesions may exert distinct biological effects (5,31,32). Although precise, traceable homology-directed repair (HDR)-based gene editing would provide an ideal model to study these variants, low HDR efficiencies in quiescent HSCs and challenges in maintaining stemness *ex vivo* limit this approach. Recent advances in human *ex vivo* HSC expansion might enable clonal selection and expansion without loss of self-renewal capacity, however, this method has proven more challenging for BM-derived HSPCs than for those from umbilical cord (51). We therefore relied on efficient, cell-cycle-independent gene editing to eliminate CEBPA-p42 and concomitantly express CEBPA-p30, functionally mimicking *CEBPA^N/N^* and *CEBPA^N/C^* patient cells. While *CEBPA^C^* mutations might generate a partially functional CEBPA-p42 protein, potentially explaining the favorable prognosis of these patients, our *CEBPA^C/C^* cells exhibited no myeloproliferative phenotype and showed no selective outgrowth of the in-frame mutant clones. Instead, their phenotype resembled *Cebpa* KO and *Cebpa^C/C^* mouse models, which exhibit impaired granulocytic/monocytic differentiation while retaining erythroid and megakaryocytic potential (7–9). Notably, whereas *Cebpa* KO mice do not develop leukemia and *Cebpa^C/C^* mice predominantly progress to erythroid leukemias, *Cebpa^N/N^* and *Cebpa^N/C^* models show that loss of CEBPA-p42 with concomitant CEBPA-p30 expression supports myeloid lineage commitment and drives granulocytic leukemias (7–9). Although the *CEBPA^N/N^* mutations generated in this study resulted in pure CEBPA-p30-expressing clones, distinguishing direct CEBPA-p30-driven mechanisms from those arising from loss of CEBPA-p42 remains challenging. For example, the shift from cDC3 to cDC2-like cells could reflect loss of CEBPA-p42, which plays a key role in guiding lineage fate of common DC progenitors, but could also be a consequence of CEBPA-p30 expression changing lineage fate (52).

While the focus of this study is CEBPA-p30-driven AML, our data also provide insights into TET2 and WT1 biology in HSPCs. ScRNA-seq revealed a highly inflammatory population of mature *TET2*-mutant granulocytic cells, consistent with recent work on excessive neutrophil extracellular trap formation by TET2-deficient, low-granule neutrophils (Supplementary Fig. 6) (45). As wild-type cells within the same individual may be more sensitive to this TET2 loss-induced inflammatory microenvironment, this could contribute to clonal expansion of TET2-deficient cells as observed in clonal hematopoiesis (53). Similarly, introduction of frameshift mutations in *WT1* exon 7 enhanced engraftment specifically in NSG mouse bone marrow, potentially driven by increased inflammation or altered metabolic processes (Fig. 5H and Supplementary Fig. 2A-B, 7H-I).

Rebalancing GATA2 levels in CEBPA-p30 cells, either through *Gata2* mutations or via TET2-driven modulation of the *Gata2* distal hematopoietic enhancer, has been identified as an important mechanism in CEBPA-p30 AML biology (12). In addition, studies have shown that *WT1* mutations can impair TET2 function through direct physical interaction, substantiated by the mutual exclusivity between *TET2* and *WT1* mutations in AML (54,55). Consistent with this, an isolated *CEBPA*/*WT1*-mutant AML subclone exhibited increased GATA2 binding at subclone-specific accessible regions, further highlighting the potential role of GATA2 in CEBPA-p30 AML (19). These data suggest that *TET2, WT1,* and *GATA2* mutations converge on common regulatory pathways in CEBPA-p30 AML. Nevertheless, clinical data indicate diverse outcomes depending on the specific co-mutations within *CEBPA*-mutant patient cohorts, ranging from favorable (*GATA2* mutations) to poor (*TET2* mutations) overall survival (1), suggesting additional mechanisms are at play.

In summary, our data demonstrate the feasibility of generating reproducible human AML models through gene editing of primary BM-derived HSPCs, overcoming patient-to-patient heterogeneity. These models represent a valuable alternative for studying favorable-risk leukemias, such as *CEBPA*-mutant AML, which are typically difficult to engraft. We identify elevated cholesterol biosynthesis as a hallmark of CEBPA-p30-driven AML. Pharmacological inhibition of cholesterol biosynthesis enhances chemosensitivity of these leukemias, suggesting that targeting cholesterol metabolism may offer a promising therapeutic strategy for *CEBPA*-mutant AML patients who fail to achieve complete remission due to suboptimal chemotherapy response.

## Acknowledgements

We would like to thank Rajesh Somasundaram for his help at the Flow Cytometry Facility, Rasmus Borring Klitgaard for his help at the Flow Cytometry & Single Cell Core Facility at the University of Copenhagen, and all animal technicians at the Animal Core Facility at the Biocenter, especially Emil Qvortrup Dyring. This work was supported by funding from Rigshospitalets Forskningspulje, Dansk Kræftforskningsfond, Dagmar Marshalls Fond, Anders Hasselbalchs Fond, Fabrikant Einar Willumsens Mindelegat granted to S.U.S., The Lundbeck Foundation (R322-2019-2576) and The Danish Cancer Society (R340-A19670) granted to B.d.B., and The Danish Cancer Society (R352-A20595) granted to B.T.P. This work has been performed in the context of the Danish Research Center for Precision Medicine in Blood Cancers funded by the Danish Cancer Society (R223-A13071) and Greater Copenhagen Health Science Partners.

## Author Contributions

Conceptualization: S.U.S., B.d.B., and B.T.P.; Data curation: S.U.S., Y.L., P.A.F., and B.d.B.; Investigation: S.U.S., Y.L., P.A.F., S.G., M.B.S., J.S., S.P., N.S.M, and B.d.B.; Visualization: S.U.S., Y.L., and B.d.B.; Methodology: S.U.S., P.A.F., N.S.M., E.M.S., R.O.B., and B.d.B.; Resources: E.M.S., K.T.M., K.G., and R.O.B.; Supervision: B.d.B. and B.T.P. Writing – original draft: S.U.S. and B.d.B.; Writing – review & editing: S.U.S, Y.L., P.A.F., S.G., M.B.S., J.S., S.P., N.S.M., E.M.S., K.T.M., K.G., R.O.B., B.d.B., and B.T.P.; Funding acquisition: S.U.S., B.d.B. and B.T.P.

## Methods

### Human sample collection, HSPC isolation and expansion

BM samples were collected from 20–30-year-old healthy donors in accordance with the standard protocol of the Department of Hematology, Rigshospitalet. Written informed consent was obtained prior to sample collection, in compliance with the Declaration of Helsinki and under a protocol approved by the Danish National Ethics Committee (approval number 1705391). BM was aspirated from the posterior iliac crest and transferred to heparin-containing tubes, mixed, and placed on ice. BM cells were processed within 2 hr after collection by diluting 20mL of aspirate 1:1 with PBS containing 2 mM EDTA (Invitrogen) and 0.5% bovine serum albumin Fraction V (Roche) (PBS-mix), carefully layered over room-temperature (RT) Lymphoprep (ProteoGenix), and centrifuged at 800 × g (acceleration 9, deceleration 0) for 30 min at 20 °C. The mononuclear cell layer was collected and washed twice with PBS-mix: first at 300 × g for 10 min to remove residual Lymphoprep and debris, then at 200 × g for 10 min to reduce platelet contamination. CD34^+^ cells were then enriched using CD34 MicroBeads and LS columns on a MACS Separator (all Miltenyi Biotec) following manufacturer’s protocol. Collected HSPCs were expanded in non-tissue culture treated 24-well plates containing StemSpan (Stem Cell Technologies) supplemented with 100 ng/mL premium grade FLT3L, IL-6, TPO, and SCF (all Miltenyi Biotec), and 35 nM UM171 (Adooq Biosciences) at a density of 1-5×10^5^ cells/mL. Cells were incubated at 37°C 5% CO_2_ for 6 days with medium replenishment every other day. For all *in vitro* cultures and *in vivo* xenotransplants, we used fresh BM HSPCs to prevent cell loss during freeze-thawing procedures.

### Cell lines

Mouse stromal-5 (MS5) cells (kindly provided by Prof. Dr. J. J. Schuringa, ACC-441) were cultured in αMEM (ThermoFisher Scientific) supplemented with 10% fetal calf serum (FCS) (HyClone) and 1% penicillin/streptomycin (pen/strep) (Gibco). MS5 cells were passaged before they reached >70% confluency, to prevent contact inhibition. U937 cells (ATCC, CRL-1593.2) were cultured in RPMI GlutaMAX medium (Gibco) supplemented with 10% FCS and 1% pen/strep. Cells were passaged every two days and plated at a density of 2×10^5^ cells/mL.

### Animal studies

All mouse experiments were approved by the Danish Animal Ethical Committee. Primary human BM cells were intravenously injected in 6- to 10-week-old NOD.Cg-Prkdcscid Il2rgtm1Wjl/SzJ (NSG) mice or NOD.Cg-Prkdcscid Il2rgtm1Wjl Tg(CMV-IL3,CSF2,KITLG)1Eav/MloySzJ (NSGS) mice (The Jackson Laboratory). Mice were sublethally irradiated with 250 cGy, 18-24hr before transplantation. Following transplantation, mice received 100mg/mL Ciprofloflaxin (Sigma-Aldrich) in their drinking water for 3 weeks. Peripheral blood (PB) chimerism levels were determined by intravenous blood collection every 3 weeks, beginning 8 weeks post-transplant. Peripheral blood (PB) was collected in EDTA-coated capillary tubes (Fisher Scientific), lysed with Pharm Lyse (BD Biosciences), and analyzed by flow cytometry. In all experiments except those related to Figure 2A-L, mice were sacrificed by cervical dislocation no later than 12 weeks post-transplant or earlier if they exhibited severe weight loss and/or decline in overall health. NSG mice displayed in Figure 2A-L were sacrificed 30 weeks post-transplant. At sacrifice, femur, tibia, and iliac bones, as well as spleen were isolated for further processing. BM cells were isolated by crushing bones with a pestle and mortar in ice-cold PBS + 3% FCS and filtered through a 40µm cell strainer (Corning). Spleen cells were crushed in ice-cold PBS + 3% FCS through a 70µm filter (Corning). Both BM and spleen cells were washed twice with ice-cold PBS + 3% FCS and analyzed by flow cytometry or resuspended in FCS + 10% DMSO and cryopreserved in liquid nitrogen.

### CRISPR/Cas9-mediated gene editing

Sequences of sgRNAs used in this study are described in Supplementary Table 1. In general, if a high editing efficiency was desired, we made use of a dual-guide strategy. In case of TET2 KO in Figure 2A-L, we initially wanted to see the clonal advantage of the TET2 KO clone and therefore chose to use only one sgRNA. For GATA2, we also used one sgRNA to model haploinsufficiency as homozygous *Gata2* loss causes HSC exhaustion (26). The AAVS1 sgRNA targets an intron of the *AAVS1* safe harbor locus (21). CEBPA sgRNA1-2 target the N-terminus, whereas CEBPA sgRNA3-4 target the C-terminal bZIP domain of CEBPA. TET2 (25) and GATA2 sgRNAs target exon 3, whereas WT1 sgRNAs target exon 7, a mutational hotspot region in *WT1*-mutant AML patients. BM-derived HSPCs were edited as described previously (56). Briefly, for electroporation of 0.5-3×10^6^ HSPCs, ribonucleoprotein (RNP) complexes were prepared by combining 3 µg Alt-R S.p. HiFi Cas9 Nuclease (Integrated DNA Technologies) and 3.18 µg sgRNA with terminal nucleotides modified with 2’-O-methyl and 3’ phosphorothioate linkages (Synthego) in a PCR tube, corresponding to a molar Cas9:sgRNA ratio of approximately 1:5. The mixture was briefly centrifuged, incubated in a thermocycler at 25 °C for 10 min to allow complex formation, and then combined according to the experimental condition. HSPCs were centrifuged at 300 x g for 10 min, resuspended in RT electroporation solution (Lonza), and 20 µL of the cell suspension was added to each PCR tube containing RNPs. The suspension was gently mixed, transferred to a 16-well electroporation strip (Lonza), and electroporated using program DZ-100 on a 4D-Nucleofector X Unit (Lonza). Immediately after electroporation, 80 µL of prewarmed StemSpan medium as described above, was added dropwise. Cells were then gently transferred to 24-well culture plates containing StemSpan medium and incubated at 37°C 5% CO_2_.

### PCR and determination of indel frequencies

Genomic DNA was isolated from a minimum of 50 and up to 0.5 ×10^6^ pelleted cells with a NucleoSpin Tissue XS Kit (Machery-Nagel) according to the manufacturer’s protocol. Targeted PCRs were performed to amplify gDNA of the edited locus (Supplementary Table 2). The *CEBPA^N^* PCR was performed using the Phusion Hot Start II DNA Polymerase kit (Thermo Scientific) with GC Buffer according to the manufacturer’s protocol. To enhance amplification of GC-rich regions, 5% DMSO and 1M betaine (Sigma-Aldrich) were added to the reaction mixture. The thermocycler protocol conditions included 2 min 98°C, 30 cycles: 30s 98°C, 20s 64°C, 30s 72°C, 5 min 72°C. In the few cases where Sanger sequencing failed, nested PCR was performed using the same conditions with nested primers (Supplementary Table 2). All other targeted PCRs were performed using the Platinum™ SuperFi II DNA Polymerase kit (Invitrogen) according to manufacturer’s protocol. The thermocycler protocol conditions included 30s 98°C, 35 cycles: 10s 98°C, 10s 60°C, 30s 72°C, 5 min 72°C. PCR product size was confirmed on a 1.5% agarose gel, and products were excised from the gel. DNA was extracted using a NucleoSpin Gel and PCR Clean-up kit (Macherey-Nagel), and products were sent for Sanger sequencing (Eurofins Genomics) with either the same forward/reverse primer as used for the targeted PCR or a nested forward/reverse primer (Supplementary Table 2). For *CEBPA^N^*PCR products, 5% DMSO was added to the sequencing reaction to improve Sanger sequencing quality. Indel analysis was performed using the ICE software (Synthego).

### MS5 co-cultures

Prior to co-culture initiation, MS5 cells were seeded onto 0.2% gelatin-coated, tissue culture-treated 6-well plates and expanded until a confluent monolayer was formed. To inhibit further proliferation, confluent MS5 cells were treated with 10 µg/mL mitomycin C (Sigma-Aldrich) for 3 hr at 37 °C, followed by three washes with PBS to remove residual mitomycin C. 24 hr post-electroporation, edited HSPCs were seeded onto the MS5 monolayer in technical triplicates. Equal cell numbers were seeded for all conditions, with 0.2-8×10^5^ edited HSPCs per well. Cells were cultured in Gartner’s medium, consisting of αMEM with nucleosides and GlutaMAX (Gibco) supplemented with 12.5% FCS (HyClone), 12.5% horse serum (Invitrogen), 1% pen/strep, 57.2 µM β-mercaptoethanol (Gibco), 1 µM hydrocortisone (Sigma-Aldrich), and 20 ng/mL premium grade SCF, TPO, IL-3, and G-CSF (all Miltenyi Biotec). Cultures were monitored regularly, and 50% of the non-adherent (supernatant) cells were harvested carefully once or twice per week, when high confluency was reached. Fresh medium was added to the remaining co-culture and harvested cells were manually counted using a Bürker-Türk and analyzed by flow cytometry and targeted sequencing.

### Colony-forming-cell (CFC) assay

24 hr post-electroporation, edited HSPCs were plated in MethoCult H4434 Classic (StemCell Technologies) supplemented with 10% IMDM (Gibco) and 1% pen/strep. Cells were plated in technical triplicates, with each replicate containing 2×10³ cells. After 14 days of incubation, colony-forming unit-granulocyte (CFU-G), CFU-monocyte (CFU-M), and burst-forming unit-erythroid (BFU-E) colonies were manually scored. For secondary and tertiary replates, cells were harvested, pooled, and replated in triplicates of 1×10^4^ cells per replicate, and colonies were scored again after 14 days.

### Flow cytometry analysis

Cells were pelleted by centrifugation at 300 × g and resuspended in PBS + 3% FCS. Fc receptors were blocked with human FcR Block and additionally with mouse FcR Block for cells derived from mouse tissues (both at 1:50, BD Biosciences). Cells were incubated with FcR blocking reagents for 5 min, followed by the addition of fluorochrome-conjugated antibodies. When antibody panels contained more than one Brilliant Violet–conjugated fluorophore, 10 µL of BD Horizon™ Brilliant Stain Buffer Plus (BD Biosciences) was added per sample. Antibodies and final staining concentrations are listed in Supplementary Table 3. Cells were stained for 30 min on ice, washed to remove unbound antibodies, and resuspended in PBS + 3% FCS supplemented with 0.2 µg/mL 7-AAD (Thermo Fisher Scientific). Samples were analyzed and/or sorted using LSRFortessa, FACSAria™ III, or FACSymphony™ S6 flow cytometers (BD Biosciences).

### Western Blot

Cells were lysed in ice-cold standard RIPA buffer supplemented with protease inhibitors for 30 min on ice. Following centrifugation at 11,000 x g for 20 min at 4°C, the supernatant containing soluble proteins was collected, diluted 1:1 in 2x Laemmli buffer supplemented with 10mM DTT and boiled for 10 min. Proteins were separated by SDS-PAGE using NuPAGE 4–12% Bis-Tris gels (Invitrogen) (1×10^5^ cells per lane) and transferred onto 0.45-μm PVDF membranes (Millipore) overnight at 4°C. For TET2 immunoblots, 0.01% SDS was included in the transfer buffer to facilitate protein transfer. Membranes were blocked for 1 hr at RT in 5% skim milk and incubated with primary antibodies overnight at 4°C. The following primary antibodies were used: CEBPA (#2295, Cell Signaling), TET2 (#A304-247A, Bethyl Laboratories), GATA2 (#4595, Cell Signaling), WT1 (CAN-R9(IHC)-56-2, Abcam), and Vinculin (#13901, Cell Signaling). Primary antibodies were detected using horseradish peroxidase-conjugated anti-rabbit secondary antibody (DAKO). Protein bands were detected using SuperSignal™ West Femto Maximum Sensitivity Substrate or SuperSignal™ West Atto Ultimate Sensitivity Substrate (Thermo Scientific).

### scRNA-seq

scRNA-seq was performed using On-Chip Multiplexing technology with the Gem-X Universal 3’ Gene Expression v4 4-plex kit according to manufacturer’s protocol (10x Genomics). Briefly, cryopreserved BM cells from each condition were thawed in 10 mL pre-warmed RPMI GlutaMAX medium (Gibco) + 20% FCS, spun down at 300 x g, resuspended in 5 mL pre-warmed RPMI GlutaMAX medium + 20% FCS supplemented with 100 µg/mL DNAse (Roche) and 5mM MgCl_2_, and incubated at 37°C for 15 min. Cells were spun down and stained for flow cytometry analysis as described in the relevant method section. For each condition, 3,000 hCD45^+^CD3^−^CD19^−^CD34^+^ and 3,000 hCD45^+^CD3^−^CD19^−^CD34^−^ cells from two individual mice were sorted sequentially into the same collection tube, resulting in 6,000 cells per population for each condition. Cells were sorted into DNA LoBind® Eppendorf tubes containing 200 µL RPMI GlutaMAX medium + 10% FCS using the FACSymphony™ S6 (BD Biosciences). After sorting, cells were placed on ice and processed on the same day. The two resulting libraries were quantified using a Qubit fluorometer (Thermo Fisher Scientific) and average fragment sizes were determined using a TapeStation (Agilent). The libraries were pooled at equimolar ratios and sequenced on a NovaSeq 6000 (Illumina) with an S1 reagent kit using paired-end reads and dual indexing.

### scRNA-seq data analysis

#### Data preprocessing, normalization, and clustering

Raw scRNA-seq data were aligned and quantified using Cell Ranger (v9.0.1; 10x Genomics) with default parameters. Downstream analyses were performed in Python using Scanpy (v1.11.1) (57). Cells with fewer than 200 detected genes, more than 10% mitochondrial gene content, or genes expressed in fewer than three cells were excluded. Doublets were identified and removed using Scrublet (v2.0.1) with the following parameters: min_counts=2, min_cells=3, min_gene_variability_pctl=85, n_prin_comps=40 (58). After quality control, a total of 30,190 cells were retained for downstream analyses. Data were normalized and log-transformed, and the top 2,000 highly variable protein-coding genes were selected for subsequent analyses. No batch effects were detected, as all samples were processed identically. Dimensionality reduction was performed using principal component analysis (PCA), followed by construction of a k-nearest neighbor graph and visualization using Uniform Manifold Approximation and Projection (UMAP). Cell clustering was performed using the Leiden algorithm with an empirically determined resolution.

#### Cell type annotation

Cell type annotation was performed using the BoneMarrowMap R package (24). To refine cell identity resolution, Leiden clustering was applied to subdivide cells into smaller clusters. Cell types were annotated based on multiple reference datasets, including the hematopoietic cell type annotations from Bandyopadhyay *et al.* (37), the *NovershternHematopoieticData* reference from the SingleR R package (v2.11.3) (59), the *Immune_All_High* reference from the Celltyplist Python package (v1.7.1) (60), and previously described dendritic cell subsets from Villani *et al.* (43). Gene signatures for leukemic stem cells (LSC), granulocyte–monocyte progenitors (GMP), erythroid, and lymphoid lineages were obtained from published studies and used to calculate cell type–specific signature scores (35,61–63). Additionally, expression of manually curated marker genes was evaluated to confirm cell identity (34–39).

#### Differential gene expression analysis of cDC lineages and CEBPA expression in HSC/MPPs

Differential gene expression analysis between cDC2-like and cDC3 cells was performed using the *FindMarkers* function in Seurat (v5.3) (64), with *ident.1 = “cDC2 like”*, *ident.2 = “cDC3”*. Genes were considered significantly differentially expressed if they met the criteria of an average log2 fold change (avg_log2FC) > 0.5 and an adjusted p-value < 0.05. For analysis of *CEBPA* expression in HSC/MPPs, RNA expression levels were visualized using boxplots generated with the ggpubr R package. Statistical comparisons across samples were performed using the Wilcoxon rank-sum test.

#### Re-clustering of the monocyte lineage

To investigate monocytic differentiation in greater detail, a subset of cells corresponding to HSC/MPP, early GMP, late GMP, pro-monocytes, and monocytes was extracted from the full dataset. Re-clustering was performed as described above using PCA-based neighborhood graph construction with *n_neighbors = 20* and *n_pcs = 40* followed by UMAP visualization. Clustering was conducted using the Leiden algorithm with a resolution of 0.6.

#### Pathway enrichment analysis of the monocytic lineage

Pathway enrichment analysis was performed for each monocyte lineage cluster using the clusterProfiler R package (65–67), with the Hallmark gene set collection as the reference dataset. First, cluster-specific upregulated genes were identified using the *FindAllMarkers* function in Seurat (v.5.3) with the parameters *only.pos = TRUE*, *min.pct = 0.25*, and *logfc.threshold = 0.5*. Next, the *compareCluster* function was used to perform enrichment analysis across clusters. Enrichment results were visualized using the *dotplot* function.

### Mass spectrometry

#### Sample preparation and processing

Cryopreserved BM cells were thawed as described in the scRNA-seq method section followed by antibody staining as described in the flow cytometry method section. Cells were processed according to a recently developed methodology performing proteomics on as few as 500 cells (68). Briefly, cells were sorted on a FACSymphony™ S6 (BD Biosciences) using a 100-micron nozzle into a twin.tec 384 LoBind plate (Eppendorf) containing 1µl of 0.2% DDM, 80mM TEAB pH 8.5 lysis buffer. After the sort, plates were briefly spun down, snap-frozen on dry ice, and boiled at 95°C for 5 min to complete the cell lysis using a Veriti thermal cycler (Applied Biosystems). Subsequently, the heated plates were allowed to cool down on ice and proteins were digested by adding 1 µL containing 10 ng Trypsin Gold (Promega) in 100 mM TEAB and incubating at 37°C overnight. The next day, each well was acidified by the addition of 1 μL 1 % (v/v) trifluoroacetic acid (TFA) and stored at −80°C until further processing. All liquid dispensing into 384-well plates was done using an I-DOT One instrument (Dispendix). All 384-well plates were thawed in the fridge and each well was loaded onto Evotip Pure according to the manufacturer’s instructions. Briefly, the tips were rinsed with 20 μl Solvent B (MS-grade ACN with 0.1% formic acid), conditioned by soaking in 2-propanol for 30 sec, equilibrated with 20 μl Solvent A (MS-grade water with 0.1% formic acid). The centrifugation steps in between were always at 700xg for 60 seconds. The tips were placed in a box filled with Solvent A to keep Evotips wet while sample loading. Each well containing 500 cells (around 4.5 µl) was loaded onto wet tips containing 15 μl solvent A, the wells were rinsed with 5 µl Buffer A* (2% ACN, 0.1% TFA) and loaded onto tips. The loaded tips were centrifuged at 700xg for 60 seconds and washed with 20 μl solvent A. Finally, tips were filled with 250 μl Solvent A, and submerged in a box with Solvent A.

#### Liquid chromatography

Chromatographic separation of peptides was conducted on an EvosepOne UHPLC system (Evosep) connected to a 15 cm Aurora Elite™ TS (Ion Opticks). All separations were carried out with the column oven set to 50°C and utilizing Evosep’s built-in 20 samples per day method.

#### Mass spectrometry

The acquisition of peptides from 500 cells was conducted using an Orbitrap Eclipse Tribrid Mass Spectrometer (Thermo Fisher Scientific), which was operated in positive mode and equipped with the FAIMS Pro interface. For this process, a single compensation voltage of −45 V was utilized, as described previously (69). The MS1 spectra were obtained using the Orbitrap, with a set resolution of 120,000 and a scan range between 400 and 1000 m/z. Additionally, the normalized automatic gain control target was set at 300%, alongside a maximum injection time of 246 ms.

### Mass-spectrometry-based proteomics data analysis

#### Raw data analysis

DIA data were processed using DIA-NN v1.9.2 (70). Raw files were searched in library-free mode with *in silico* spectral library generation enabled. Peptide identification was performed against a combined FASTA database comprising the reviewed *Homo sapiens* proteome (UniProt, November 2024). Trypsin specificity was assumed with cleavage at K/R residues, allowing up to one missed cleavage. The precursor m/z range was set to 400–800 m/z with charge states +1 to +4, and fragment ions were considered in the 200–1800 m/z range. Peptide lengths were restricted to 7–30 amino acids. N-terminal methionine excision was enabled. Oxidation of methionine was included as the only variable modification, with a maximum of one variable modification per peptide. False discovery rate was controlled at 1% at the precursor and protein levels. Quantification matrices for precursors and protein groups were exported for downstream analysis.

#### Data processing and imputation

All subsequent data processing and statistical analyses below were in R (v4.4.3). Technical replicates with low protein identification depth (< 5,000 detected proteins) were excluded, and one *GATA2* single-mutant biological replicate was removed due to low indel frequency. Following sample exclusion, the DIA-NN analysis described above was re-run, yielding a final protein abundance matrix containing 7,186 detected proteins. Sequential filtering steps were applied as follows: (i) proteins lacking gene name annotations were removed; (ii) for duplicated gene names, isoforms with the lowest detection across samples or with non-hematopoietic tissue specificity were excluded; (iii) proteins detected in < 2 technical replicates per biological replicate across all conditions were removed; and (iv) proteins detected in < 2 biological replicates across all condition were excluded. After filtering, 7,037 proteins remained. Protein abundances were log2-transformed, and technical replicates were collapsed by computing the mean abundance per biological replicate, resulting in 91.6% global data completeness. Remaining missing values were assumed to be missing not at random and were imputed using quantile regression imputation of left-censored data (QRILC) via the *impute.QRILC* function in the imputeLCMD R package (v2.1) (71), with a fixed random seed (set.seed(1)) for reproducibility. All downstream analyses were conducted on this log2-transformed, imputed dataset.

#### Principal component analysis

Principal component analysis (PCA) was performed using the *prcomp* function in R with centering and scaling (center = TRUE, scale = TRUE). PCA was performed on three data subsets: (i) AAVS1, CEBPA, TET2, and CEBPA+TET2, (ii) AAVS1, CEBPA, GATA2, and CEBPA+GATA2, and (iii) AAVS1, CEBPA, WT1, and CEBPA+WT1. Principal component scores were extracted and visualized as PC1 vs. PC2 scatter plots.

#### Differential expression analysis

Differential expression analysis was performed using the limma R package (v3.62.2). A single linear model including all conditions was fitted, and array weights were estimated using the *arrayWeights* function to account for sample heterogeneity. Contrasts were defined to compare each condition to the AAVS1 control, and empirical Bayes moderation was applied to stabilize variance estimates across proteins. For each contrast, proteins were retained only if they contained at least two measured (non-imputed) values in the test condition and/or the AAVS1 control. Proteins meeting this requirement were considered differentially expressed if they exhibited |log2 fold change| > log2(1.3) and p-value < 0.05. Differentially expressed proteins were categorized according to directionality, annotated according by condition, and visualized using pheatmap (v1.0.13). Pathway enrichment analysis was performed using the *compareCluster* function from the clusterProfiler R package (v4.14.6) with ReactomePA (v1.50.0), and results were visualized using the *dotplot* function from the same package.

### Drug screening

Prior to cell plating, cytarabine (AraC) (MedChemExpress), simvastatin (MedChemExpress), and DMSO were dispensed into 384-well polypropylene V-bottom plates (Greiner) using an Echo 550 Acoustic Liquid Handler (Beckman Coulter). Compound placement was randomized across wells to minimize potential positional effects. On the day of treatment, supernatant cells were harvested from MS5 co-cultures, pooled by condition, manually counted, and resuspended in fresh Gartner’s media supplemented with cytokines to achieve equal cell concentrations across conditions. Then, 20 µL of cell suspension containing 13,000–15,000 cells was transferred in technical triplicates to each well of the compound plates. The plates were briefly centrifuged at 90 x g and incubated at 37°C 5% CO_2_ for 96 hr. Following incubation, antibodies and buffers were added to each 20 µL well using an Echo 525 acoustic liquid handler (Beckman Coulter), including DRAQ7™ (BioLegend) and anti-human CD45 PE-Cy7 (clone HI30, BioLegend). Cells were then incubated in the dark at RT for 1.5 hr prior to analysis on an iQue® 3 high-throughput screening cytometer (Sartorius).

### Statistics

For all flow cytometry data, CFC assay, and the drug screens, significance was evaluated by a one-way ANOVA followed by a Šídák’s multiple comparisons test using Graphpad Prism 10. Statistics for the scRNA-seq and mass spectrometry can be found in the relevant method sections.

---

**Supplementary Figure 1.**
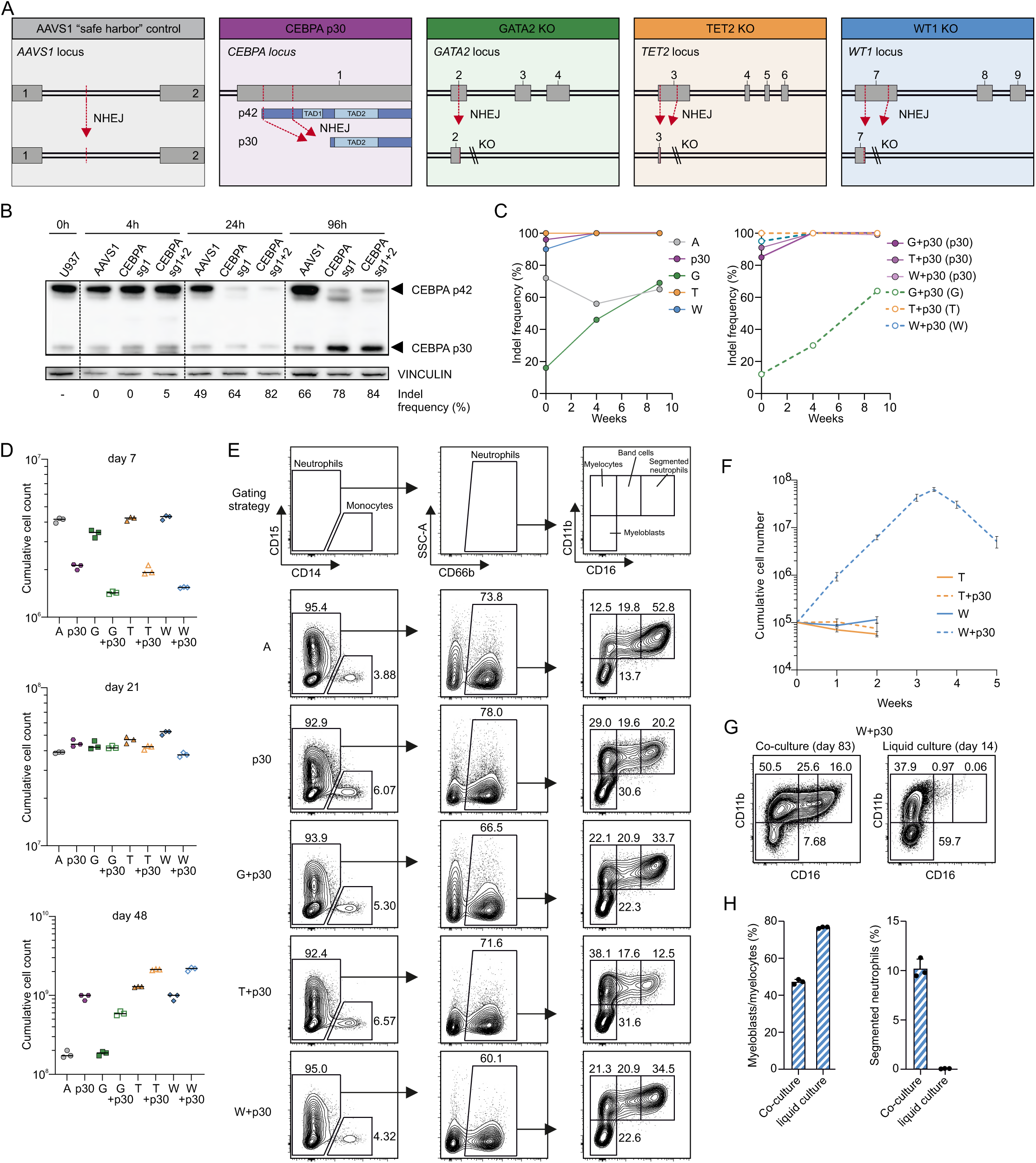
Loss of TET2 or WT1 accelerate CEBPA-p30-driven granulocytic differentiation block. **(A)** Schematic showing sgRNA binding sites in each condition (21,22). bZIP; Basic Leucine Zipper, NHEJ; non-homologous enjoining, TAD; transactivation domain. **(B)** CEBPA-p42 and CEBPA-p30 expression in U937 cells 4 hours (hr), 24 hr, and 96 hr after electroporation. Vinculin was used as loading control. Insertion/deletion (indel) frequency of the relevant gene are indicated below each lane. **(C)** Indel frequency of single-mutant (left) and double-mutant (right) bone marrow (BM) hematopoietic stem/progenitor cells (HSPCs) co-cultured on mouse stromal-5 (MS5) cells as shown in Figure 1F. In case of double-mutants, the gene between brackets is shown. From here on: A; AAVS1, p30; CEBPA-p30, G; GATA2 knockdown (KD), T; TET2 knockout (KO), and W; WT1 KO. **(D)** Cumulative cell counts of edited HSPCs grown on a stromal layer of MS5 cells at days 7, 21, and 48. Lines indicate mean of technical triplicates. **(E)** Representative gating strategy for neutrophilic differentiation in supernatant cells harvested at day 15 of co-culture. **(F)** Cumulative cell counts of supernatant cells harvested at day 69 from co-cultures shown in Figure 1F. A portion of these cells were subsequently grown in liquid culture. Error bars represent mean ±SD of technical triplicates. **(G-H)** Representative gating strategy **(G)** and percentage of myeloblasts/myelocytes (left) and segmented neutrophils (right) **(H)** in W+p30 cells harvested from co-cultures (day 83) and liquid cultures (day 14). Bar plots represent mean ± SD. *p<0.05, **p<0.01, ***p<0.001.

**Supplementary Figure 2.**
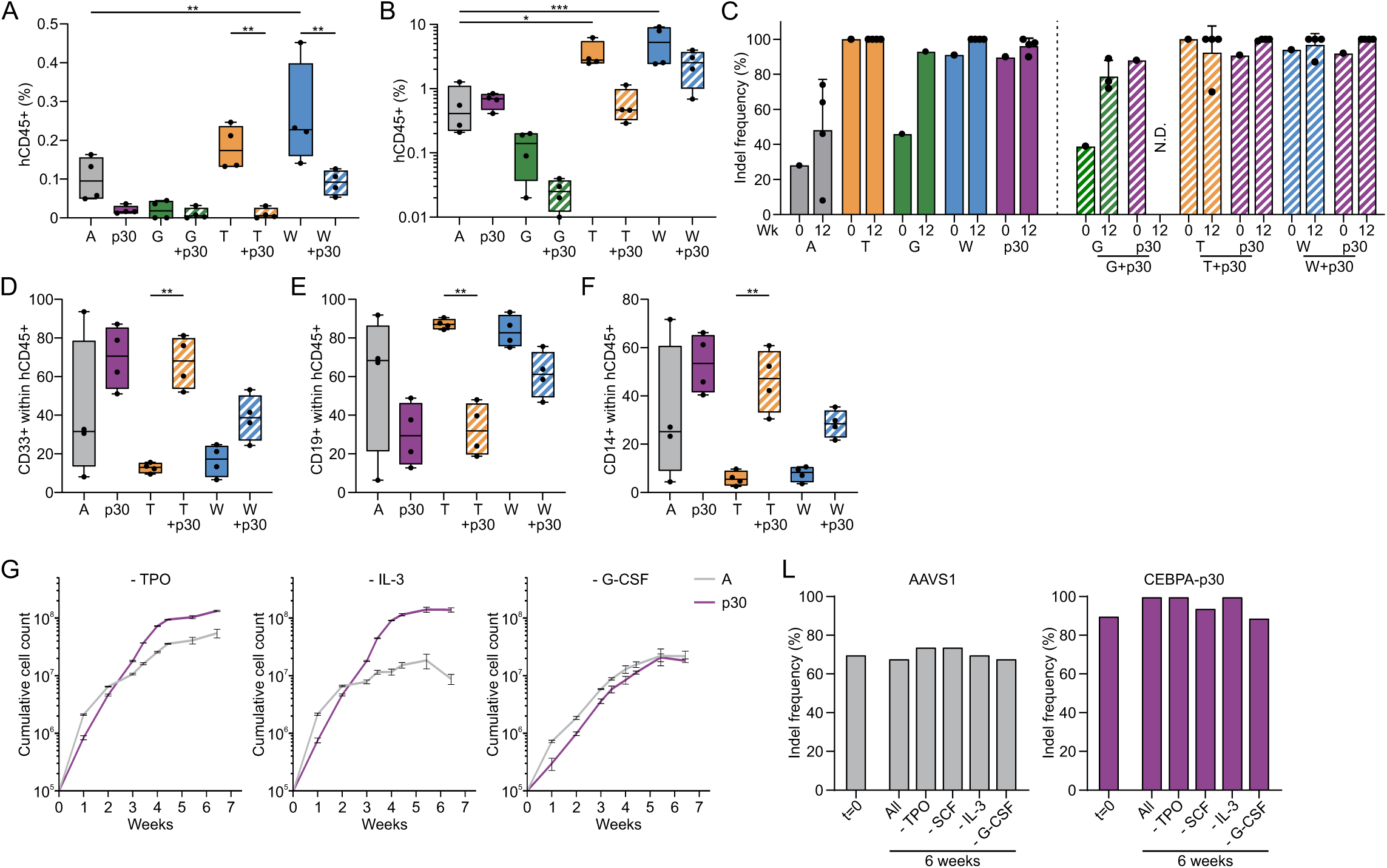
CEBPA-p30 cells rely on human cytokines and interaction with the BM microenvironment. **(A-B)** Percentage of human CD45^+^ (hCD45^+^) cells in peripheral blood (PB) at week 8 **(A)** and BM at sacrifice **(B)** in NSG mice. **(C)** Indel frequency at time of transplantation (week (wk) 0) and in BM hCD45^+^CD3^−^CD19^−^ cells at sacrifice (wk 12). **(D-F)** Percentage of CD33^+^ **(D)**, CD19^+^ **(E)**, and CD14^+^ **(F)** cells within hCD45^+^ cells in BM. **(G)** Cumulative cell counts of edited HSPCs grown on MS5 cells in medium lacking either thrombopoietin (TPO), interleukin-3 (IL-3) or granulocyte-colony stimulating factor (G-CSF). Mean ±SD of technical triplicates. **(L)** Indel frequency of A (left) and p30 (right) at plating (t=0) and after 6 weeks in co-culture with different cytokines cocktails. Boxplots show median (line), interquartile range (box), and minimum/maximum values (whiskers). N.D.; not determined. *p<0.05, **p<0.01, ***p<0.001.

**Supplementary Figure 3.**
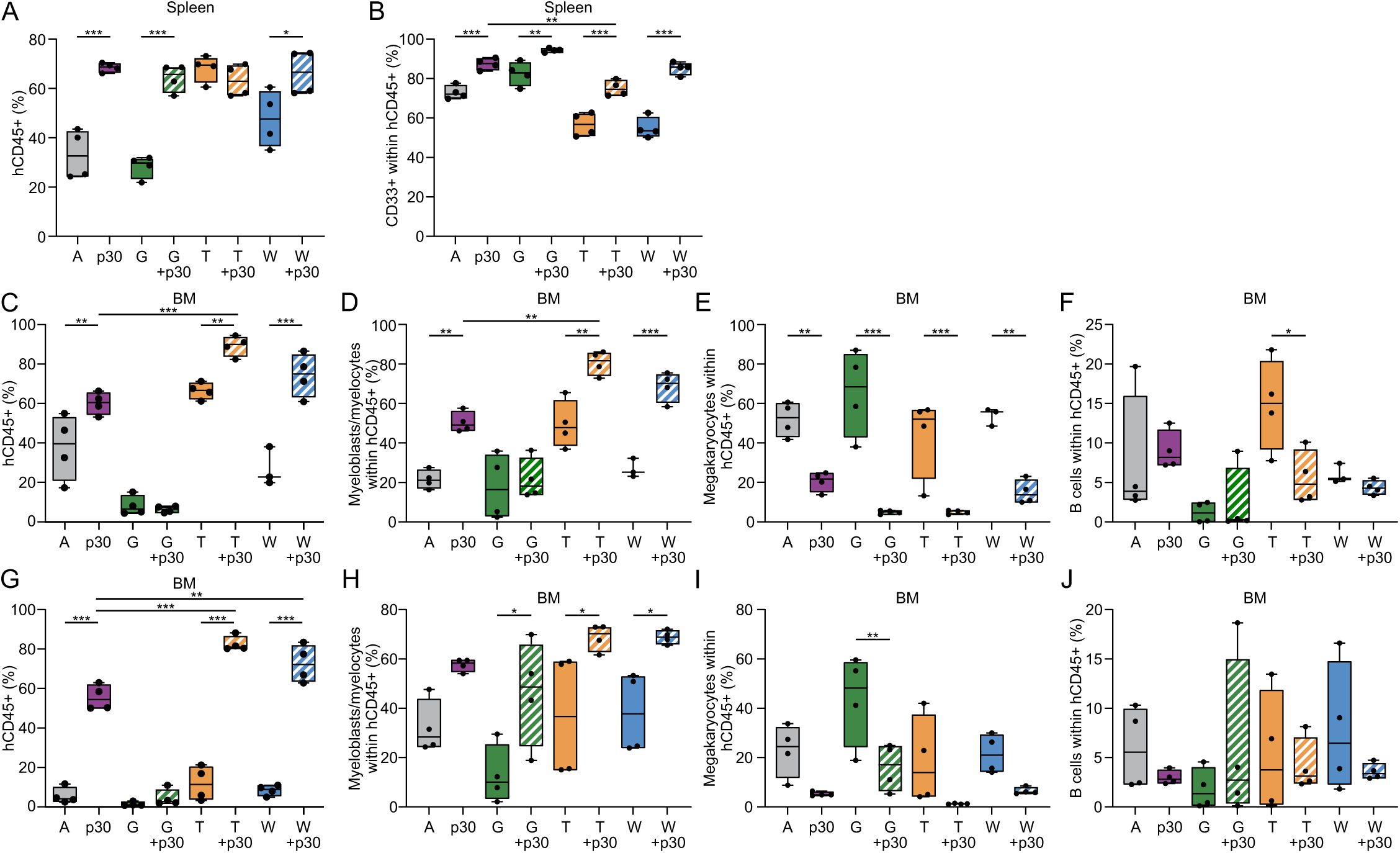
CEBPA-p30 combined with TET2 or WT1 loss induces lethal AML in NSGS mice. **(A-B)** Percentage of hCD45^+^ cells **(A)** and CD33^+^ cells within the hCD45^+^ compartment **(B)** in the spleen at sacrifice. Related to Figure 3F. **(C)** Percentage of hCD45^+^ cells in the BM of NSGS mice following gene editing and transplantation of BM-derived CD34^+^ HSPCs from an independent donor. **(D-F)** Percentage of myeloblasts/myelocytes (CD66b^+^CD16^−^) **(D)**, megakaryocytic cells (CD41a^+^) **(E)**, and B cells (CD19^+^) **(F)** within BM-derived hCD45^+^ cells at sacrifice. **(G)** Percentage of hCD45^+^ cells in the BM of NSGS mice following gene editing and transplantation of BM-derived CD34^+^ HSPCs from another independent donor. **(H-J)** Myeloblasts/myelocytes (CD66b^+^CD16^−^) **(H)**, megakaryocytic cells (CD41a^+^CD117^+^) **(I)**, and B cells (CD19^+^) **(J)** within BM-derived hCD45^+^ cells at sacrifice. Box plots show median (line), interquartile range (box), and minimum/maximum values (whiskers). *p<0.05, **p<0.01, ***p<0.001.

**Supplementary Figure 4.**
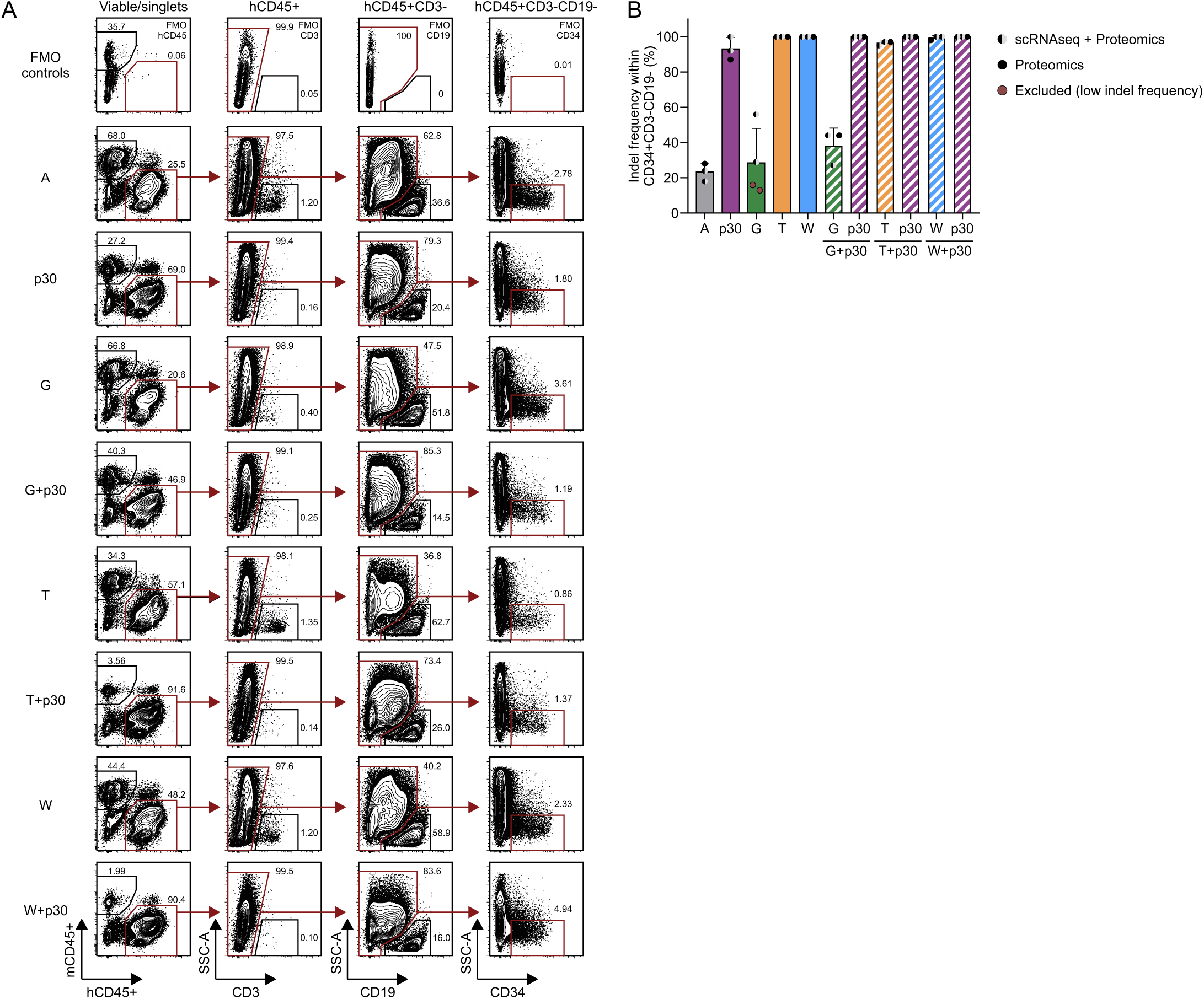
Gating strategy and gene editing efficiencies of cells used for single-cell RNA sequencing (scRNA-seq) and low-input proteomics. **(A)** Representative gating strategy used to sort BM-derived hCD45+CD3-CD19-CD34+ HSPCs (scRNA-seq and proteomics) and hCD45+CD3-CD19-cells (scRNA-seq) from NSGS mice for each experimental condition. FMO; fluorescence minus one. **(B)** Indel frequency within BM-derived hCD45+CD34+CD3-CD19-HSPCs from NSGS mice. GATA2-mutant samples indicated by a red dot were excluded from analysis due to low GATA2 editing efficiency.

**Supplementary Figure 5.**
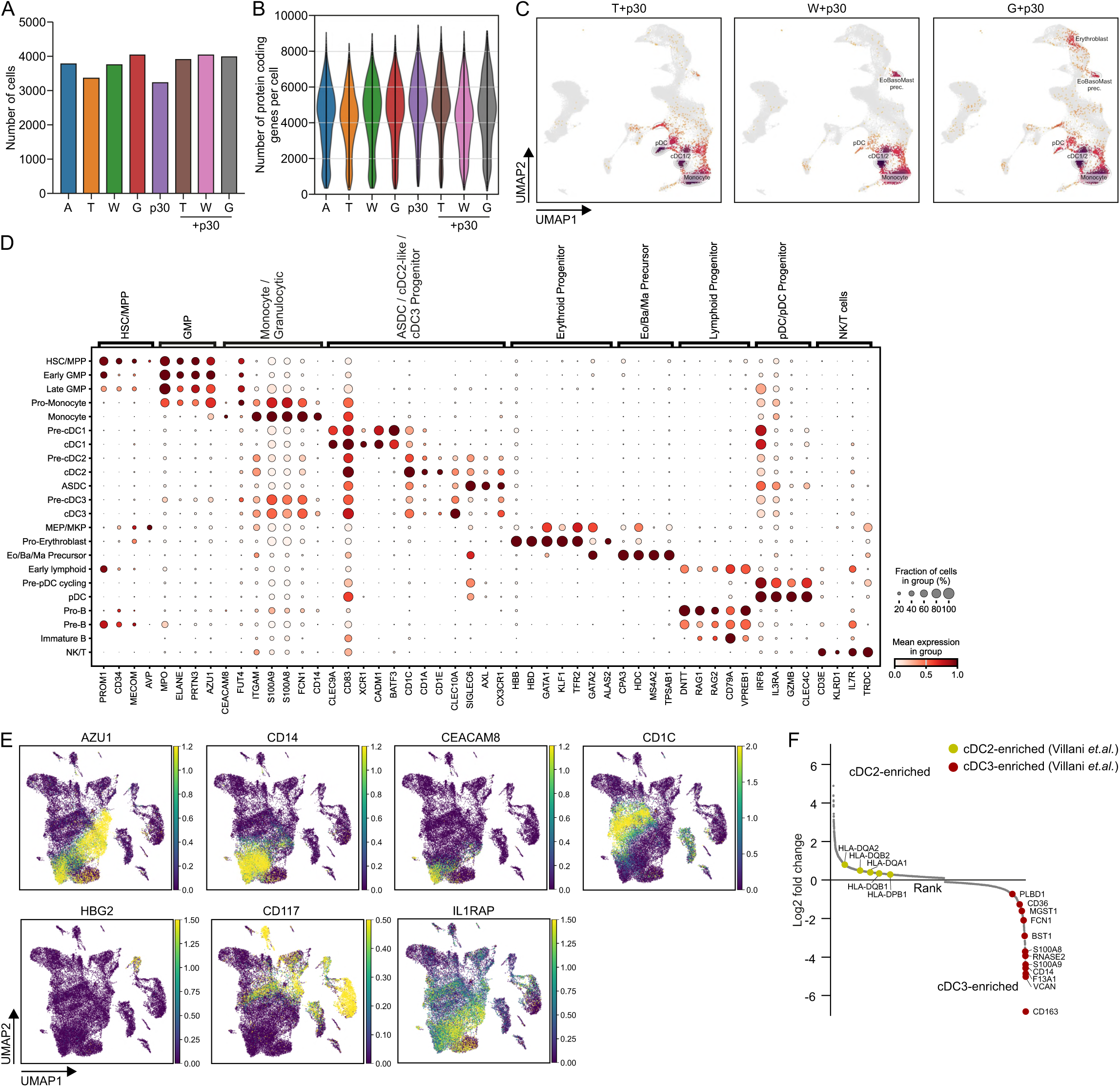
scRNA-seq confirms CEBPA-p30-driven myeloid bias and reveals a shift toward cDC2-like cells. **(A)** Number of detected cells per condition using on-chip multiplexing scRNA-seq. **(B)** Number of annotated protein coding genes per cell across condition. **(C)** Projection of CEBPA-p30 double-mutant cells onto a healthy reference UMAP previously generated by Zeng *et al.*(*56*). **(D)** Dot plot of gene expression highlighting genes used to annotate the UMAP clusters shown in Figure 5B. **(E)** Expression of selected genes plotted on the reference map of all cell types. **(F)** Differential gene expression within the cDC2 and cDC3 clusters. Genes distinguishing cDC2 (yellow) and cDC3 (red), as described by Villani *et al.* (29) are highlighted.

**Supplementary Figure 6.**
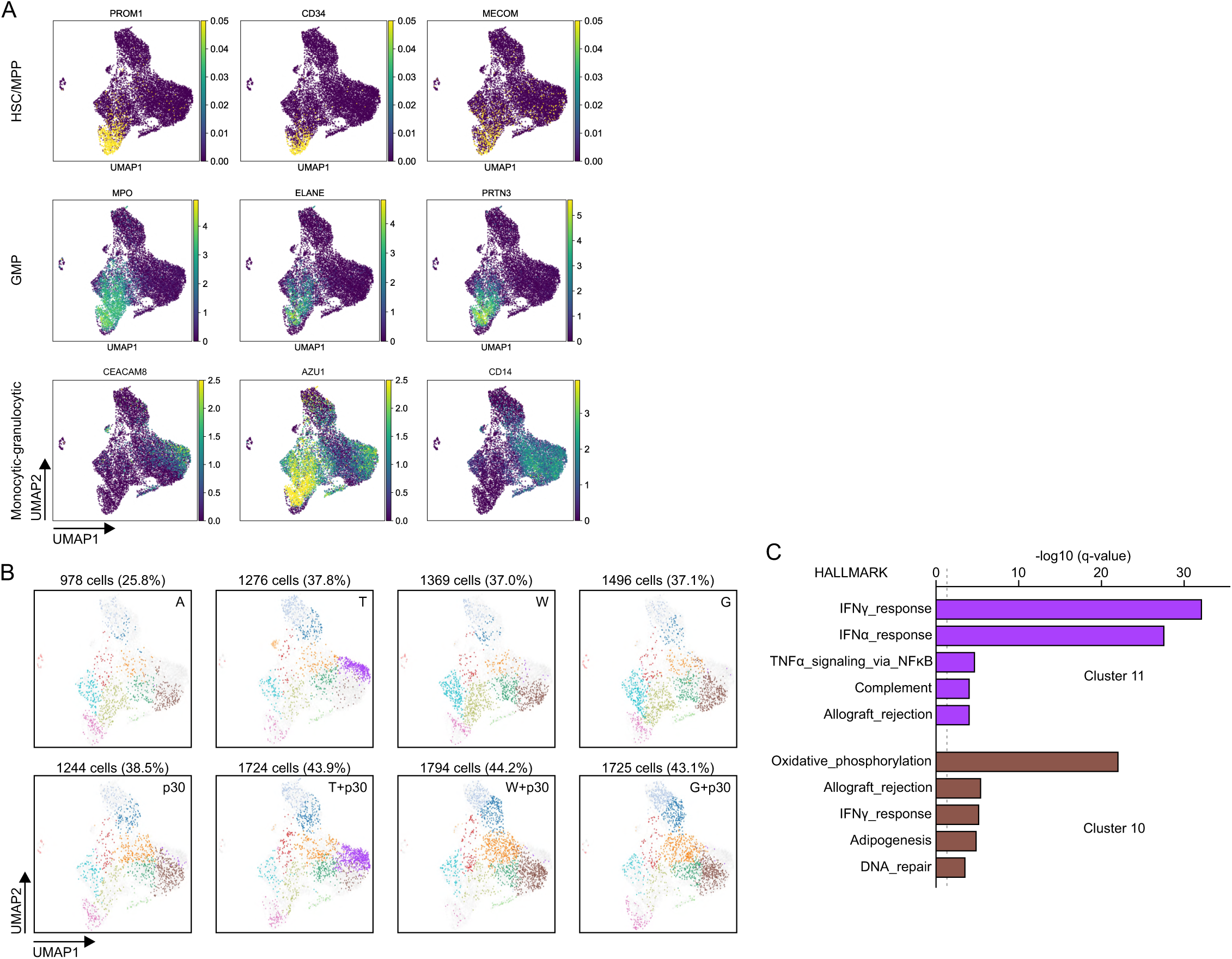
TET2 loss results in a subset of highly inflamed granulocytes. **(A)** Examples of genes that contribute to defining the HSC/MPP, GMP, and monocytic/granulocytic compartments. **(B)** UMAPs of cells from individual conditions projected onto the common UMAP shown in Figure 5I. **(C)** Pathway enrichment analysis of genes expressed in cluster 11 (purple) and cluster 10 (brown). Related to Figure 5I-J.

**Supplementary Figure 7.**
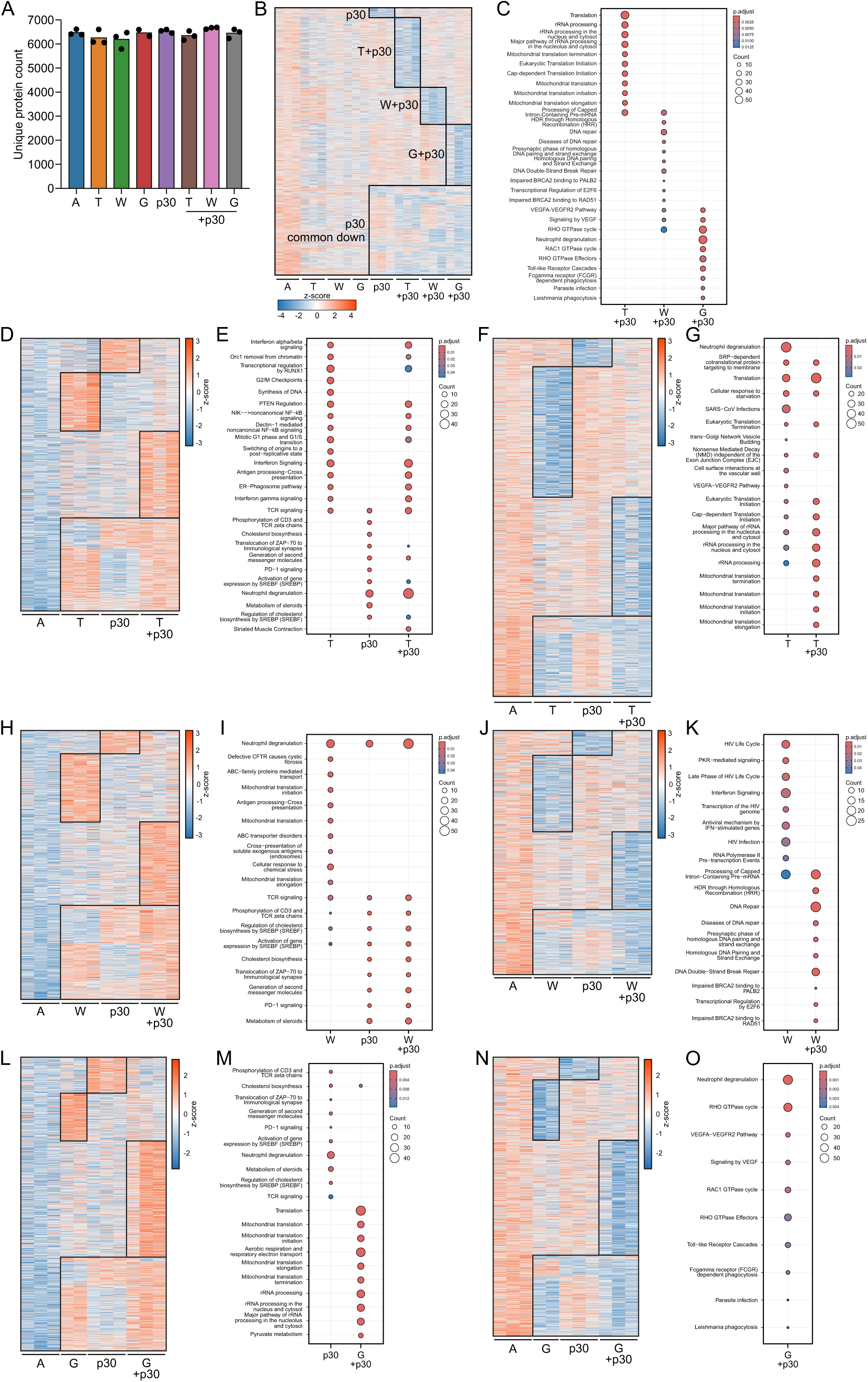
Unique and shared biological processes driven by single- and/or double-mutant HSPCs. **(A)** Number of unique proteins identified in technical replicates for each condition. **(B-C)** Proteins downregulated (log2FC < −log2(1.3), p<0.05) in CEBPA-p30 single- and/or double-mutants relative to AAVS1 HSPCs **(B)** and the corresponding enriched Reactome pathways **(C)**. **(D-G)** Upregulated **(D)** and downregulated **(F)** proteins and related Reactome pathways **(E and G)** in T, p30, and T+p30 cells relative to AAVS1 HSPCs. **(H-K)** Upregulated **(H)** and downregulated **(J)** proteins and related Reactome pathways **(I and K)** in W, p30, and W+p30 cells relative to AAVS1 HSPCs. **(L-O)** Upregulated **(L)** and downregulated **(N)** proteins and related Reactome pathways **(M and O)** in G, p30, and G+p30 cells relative to AAVS1 HSPCs.

**Supplementary Figure 8.**
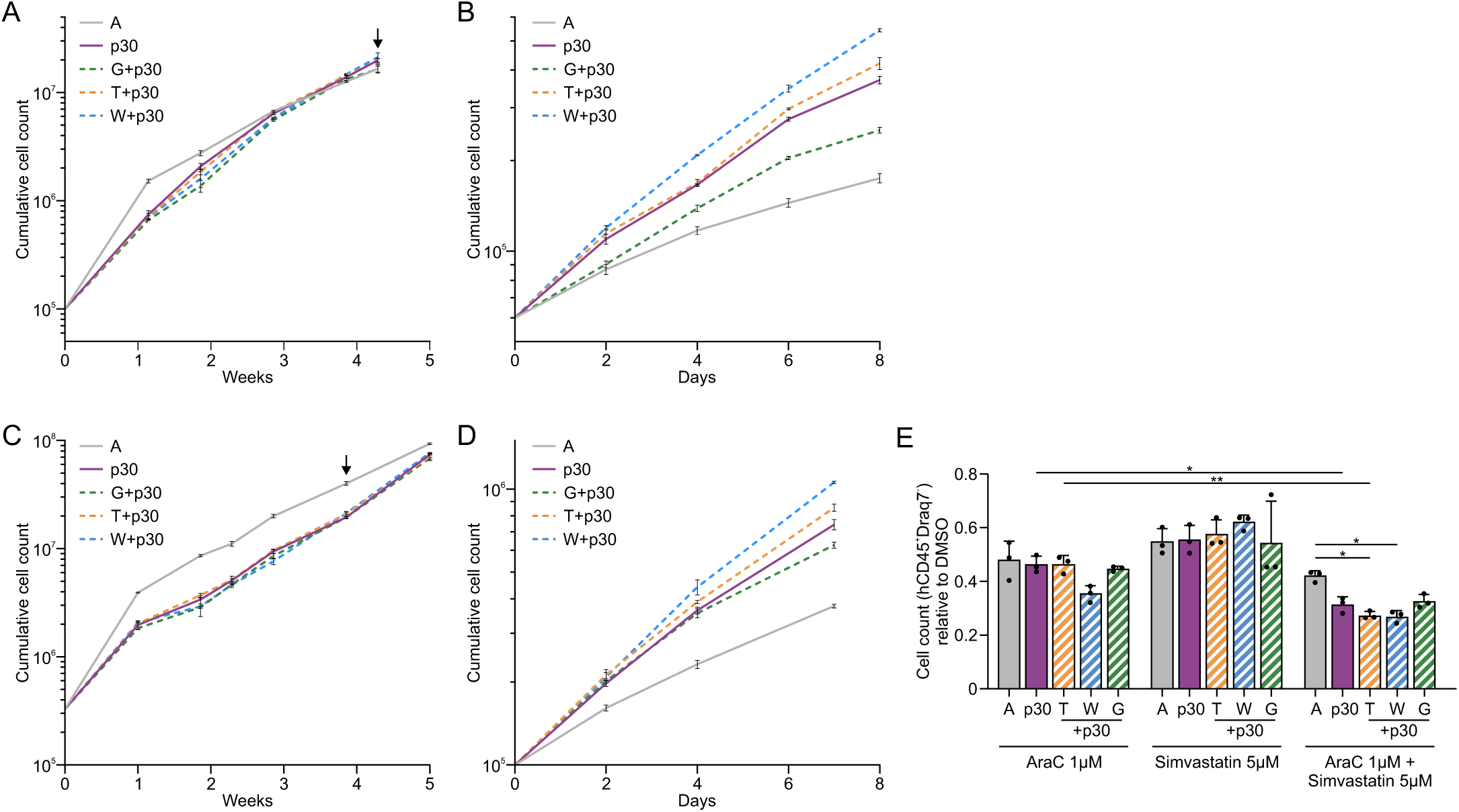
Targeting cholesterol biosynthesis increases chemosensitivity of CEBPA-p30 cells. **(A-B)** Cumulative cell counts of edited HSPCs grown on a stromal layer of MS5 cells **(A)**. Supernatant cells were harvested after 30 days (arrow) and either used for a drug sensitivity assay (Figure 6G) or grown in liquid culture for up to 8 days **(B)**. **(C-E)** BM-derived CD34^+^ HSPCs from an independent donor were subjected to gene editing, expansion, MS5 co-culture, and drug sensitivity analysis as in (A-B) and Figure 6G. Cumulative cell counts of edited HSPCs grown on a stromal layer of MS5 cells **(C)**. Supernatant cells were harvested after 28 days (arrow) and either grown in liquid culture for up to 7 days **(D)** or directly used for a drug sensitivity assay with viable (hCD45^+^DRAQ7^−^) cell numbers read out after 96 hr **(E)**. Error bars indicate mean ± SD of technical triplicates. *p<0.05, **p<0.01, ***p<0.001.

**Supplementary Table 1.**
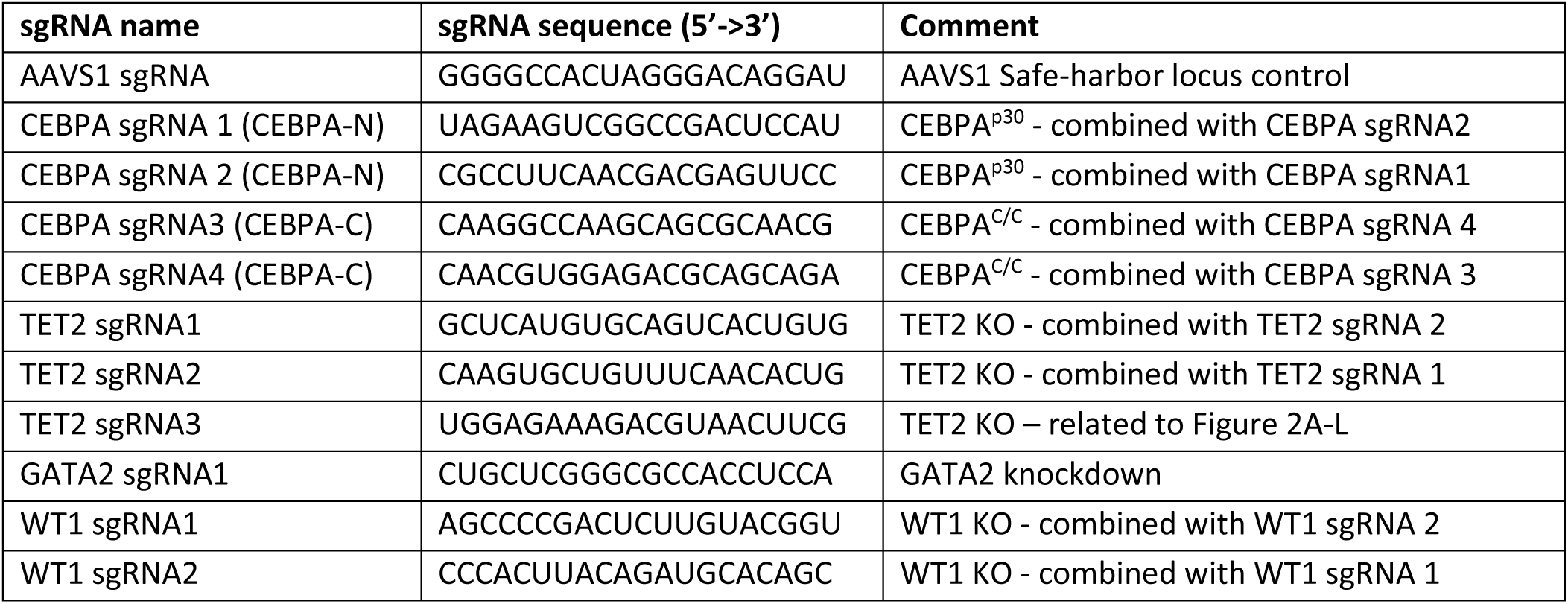
Sequences of sgRNAs used in this study.

**Supplementary Table 2.**
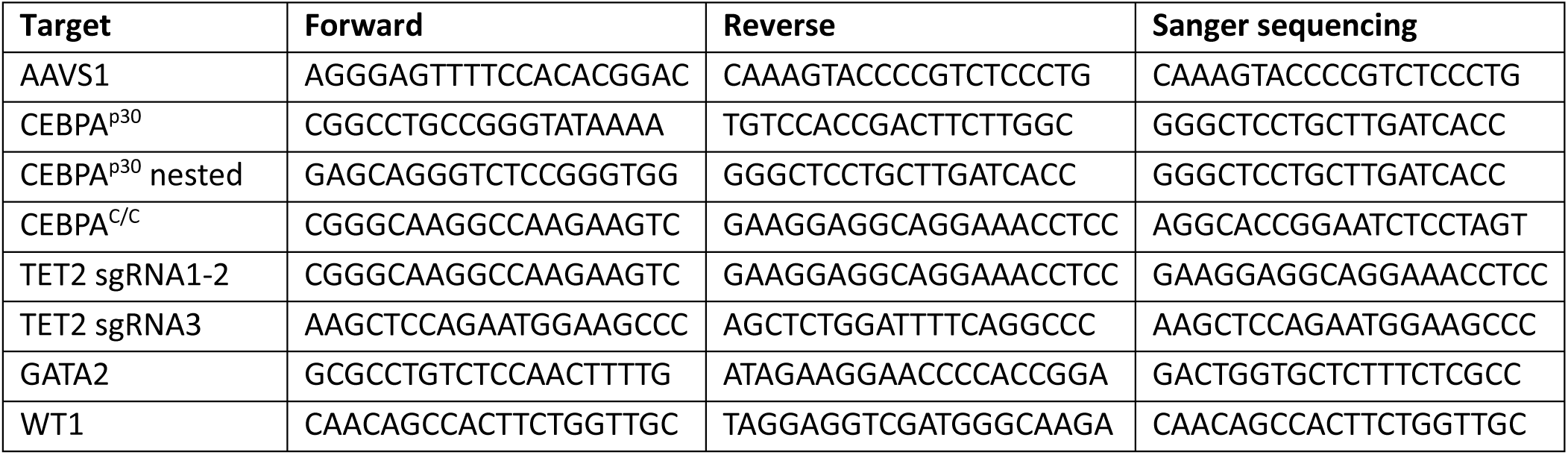
Oligonucleotides used for PCR and sanger sequencing in this study.

**Supplementary Table 3.** Antibodies for flow cytometry used in this study.

| Marker | Fluorophore | Clone | Company |
| --- | --- | --- | --- |
| 7AAD |  |  | ThermoFisher |
| CD117 | FITC | 104D2 | Invitrogen |
| CD11b | APC-Cy7 | ICRF44 | BD Pharmingen |
| CD11b | PE | ICRF44 | Invitrogen |
| CD14 | BV786 | MΦP9 | BD Biosciences |
| CD14 | PECy7 | HCD14 | BioLegend |
| CD15 | PE | W6D3 | BD Pharmingen |
| CD15 | FITC | MMA | BD Biosciences |
| CD16 | BV605 | 3G8 | BD Biosciences |
| CD16 | BV510 | 3G8 | BD Biosciences |
| CD19 | BV786 | 2H7 | BD Biosciences |
| CD19 | PE-Cy7 | HIB19 | BD Pharmingen |
| CD3 | BV650 | SK7 | BD Biosciences |
| CD33 | BV605 | P67.6 | BD Biosciences |
| CD33 | PE | WM53 | BD Pharmingen |
| CD34 | PerCp-Cy5.5 | 8G12 | BD Biosciences |
| CD34 | APC | 581 | BD Biosciences |
| CD38 | APC-H7 | HB7 | BD Biosciences |
| CD41a | PE-Cy7 | HIP8 | BD Pharmingen |
| CD45RA | BV510 | HI100 | BD Biosciences |
| CD66b | APC | G10F5 | BioLegend |
| CD71 | BV711 | M-A712 | BD Biosciences |
| CD71 | PE | M-A712 | BD Biosciences |
| DRAQ7 |  |  | BioLegend |
| hCD45 | BV421 | HI30 | BD Biosciences |
| hCD45 | PE-Cy7 | HI30 | BioLegend |
| mCD45 | BV711 | 30-F11 | BD Biosciences |
| mCD45 | FITC | UCHT2 | BD Biosciences |

**Supplementary Data 1**

Raw scRNA-seq data files will be made publicly available through European Genome-phenome

Archive (EGA). Filtered count matrices and scripts will be made publicly available through Zenodo.

**Supplementary Data 2**

Raw proteome data will be made publicly available through EGA.

Count matrices and scripts will be made publicly available through Zenodo.

